# SARS-CoV-2 neutralizing human recombinant antibodies selected from pre-pandemic healthy donors binding at RBD-ACE2 interface

**DOI:** 10.1101/2020.06.05.135921

**Authors:** Federico Bertoglio, Doris Meier, Nora Langreder, Stephan Steinke, Ulfert Rand, Luca Simonelli, Philip Alexander Heine, Rico Ballmann, Kai-Thomas Schneider, Kristian Daniel Ralph Roth, Maximilian Ruschig, Peggy Riese, Kathrin Eschke, Yeonsu Kim, Dorina Schäckermann, Mattia Pedotti, Philipp Kuhn, Susanne Zock-Emmenthal, Johannes Wöhrle, Normann Kilb, Tobias Herz, Marlies Becker, Martina Grasshoff, Esther Veronika Wenzel, Giulio Russo, Andrea Kröger, Linda Brunotte, Stephan Ludwig, Viola Fühner, Stefan Daniel Krämer, Stefan Dübel, Luca Varani, Günter Roth, Luka Čičin-Šain, Maren Schubert, Michael Hust

## Abstract

COVID-19 is a severe acute respiratory disease caused by SARS-CoV-2, a novel betacoronavirus discovered in December 2019 and closely related to the SARS coronavirus (CoV). Both viruses use the human ACE2 receptor for cell entry, recognizing it with the Receptor Binding Domain (RBD) of the S1 subunit of the viral spike (S) protein. The S2 domain mediates viral fusion with the host cell membrane. Experience with SARS and MERS coronaviruses has shown that potent monoclonal neutralizing antibodies against the RBD can inhibit the interaction with the virus cellular receptor (ACE2 for SARS) and block the virus cell entry. Assuming that a similar strategy would be successful against SARS-CoV-2, we used phage display to select from the human naïve universal antibody gene libraries HAL9/10 anti-SARS-CoV-2 spike antibodies capable of inhibiting interaction with ACE2. 309 unique fully human antibodies against S1 were identified. 17 showed more than 75% inhibition of spike binding to cells expressing ACE2 in the scFv-Fc format, assessed by flow cytometry and several antibodies showed even an 50% inhibition at a molar ratio of the antibody to spike protein or RBD of 1:1. All 17 scFv-Fc were able to bind the isolated RBD, four of them with sub-nanomolar EC50. Furthermore, these scFv-Fc neutralized active SARS-CoV-2 virus infection of VeroE6 cells. In a final step, the antibodies neutralizing best as scFv-Fc were converted into the IgG format. The antibody STE73-2E9 showed neutralization of active SARS-CoV-2 with an IC50 0.43 nM and is binding to the ACE2-RBD interface. Universal libraries from healthy human donors offer the advantage that antibodies can be generated quickly and independent from the availability of material from recovered patients in a pandemic situation.

## Main text

In 2015 Menachery et al. presciently wrote: “Our work suggests a potential risk of SARS-CoV re-emergence from viruses currently circulating in bat populations.” ^1^. Four years later, a novel coronavirus causing a severe pneumonia was discovered and later named SARS-CoV-2. The outbreak was initially noticed on a sea food market in Wuhan, Hubei province (China) at the end of 2019. The disease was named COVID-19 (coronavirus disease 2019) by the World Health Organization (WHO). Sequencing showed high identity to bat coronaviruses (CoV, in particular RaTG13), beta-CoV virus causing human diseases like SARS and MERS and, to a lesser extent, the seasonal CoV hCoV-OC43 and HCov-HKU1 ^2,3^. The spike (S) protein of SARS-CoV-2, as well as SARS-CoV, binds to the human zinc peptidase angiotensin-converting enzyme 2 (ACE2), which is expressed on numerous cells, including lung cells, heart, kidney and intestine cells, thus initiating virus entry into target cells. S protein consists of the N-terminal S1 subunit, which includes the receptor binding domain (RBD), and the C-terminal S2 subunit which is anchored to the viral membrane and is required for trimerization of spike itself and fusion of the virus and host membrane ^4–6^. The intracellular host enzyme furin cleaves the S protein between S1 and S1 during viral formation and the membrane bound host protease TMPRSS2 is responsible for the proteolytic activation of the S2’ site, which is necessary for conformational changes and viral entry ^7–10^.

Antibodies against the spike protein of coronaviruses are potential candidates for therapeutic development ^11^. Antibodies against the S1 subunit, especially against RBD, can potently neutralize SARS-CoV and MERS ^12–14^. Monoclonal human antibodies against SARS-CoV were also described to cross-react with SARS-CoV-2, some of them were able to neutralize SARS-CoV-2 ^15,16^. In other approaches monoclonal antibodies against SARS-CoV-2 were selected by rescreening memory B-cells from a SARS patient ^17^, selected from COVID-19 patients by single B-cell PCR ^18,19^ or using phage display ^20,21^. Human recombinant antibodies were successfully used for the treatment of other viral diseases. The antibody mAb114 ^22^ and the three antibody cocktail REGN-EB3 ^23^ showed a good efficiency in clinical trials against Ebola virus ^24^. The antibody palivizumab is EMA/FDA approved for treatment of a severe respiratory infection of infants caused by the respiratory syncytial virus (RSV) ^25,26^ and could be used as a guideline to develop therapeutic antibodies against SARS-CoV-2.

Antibody phage display is a powerful tool to generate human antibodies against infectious diseases ^27^. We successfully used this technology to develop *in vivo* protective antibodies against Venezuelan encephalitis virus ^28^, Western-equine encephalitis ^29,30^, Marburg virus ^31^ and Ebola Sudan virus ^32^. In this work, we generated human recombinant antibodies against the spike proteins of SARS-CoV-2 from a universal, human naive antibody gene library that was constructed before the emergence of SARS-CoV-2. Several scFv-Fc antibodies were identified which efficiently inhibited the binding of the spike protein to ACE2-expressing cells and blocked SARS-CoV-2 infection of VeroE6 cells. The best antibody in the IgG format is a potential candidate for the clinical development of a passive immunotherapy for therapeutic and prophylactic purposes.

## Results

### SARS CoV2 spike domains or subunits and human ACE2 were produced in insect cells and mammalian cells

SARS-CoV-2 RBD-SD1 (aa319-591) according to Wrapp et al. 2020 ^33^, S1 subunit (aa14-694), S1-S2 (aa14-1208, with proline substitutions at position 986 and 987 and “GSAS” substitution at the furin site, residues 682-685) and extracellular domain of ACE2 receptor were produced in insect cells using a plasmid based baculovirus free system ^34^ as well as in Expi293F cells. All antigens with exception of S1-S2 were produced with human IgG1 Fc part, murine IgG2a Fc part or with 6xHis tag in both expression systems. S1-S2 was only produced with 6xHis tag. The extracellular domain of ACE2 was produced with human IgG1 Fc part or mouse IgG2a in Expi293F cells and 6xHis tagged in insect cells. The yields of all produced proteins are given in Table 1. A graphical overview on all produced proteins is given in Supplementary Data 1. The expressed proteins were analyzed by size exclusion chromatography (SEC) (Supplementary Data 2).

**Table 1.** Antigen production.

|  | High Five cells |  |  | Expi293F cells |  |  |
| --- | --- | --- | --- | --- | --- | --- |
|  | Yield | Binding to ACE2 in ELISA | Binding to ACE2 on cells | Yield | Binding to ACE2 in ELISA | Binding to ACE2 on cells |
| **RBD-hFc** | 90 mg/L | yes | yes | 203 mg/L | yes | yes |
| **RBD-mFc** | 48 mg/L | yes | yes | 116 mg/L | yes | yes |
| **RBD-His** | 92 mg/L | yes | yes | 35 mg/L | yes | yes |
| **S1-hFc\*** | 7 mg/L | yes | yes | <1 mg/L | no | no |
| **S1-hFc** | 50 mg/L | yes | yes | <1 mg/L | weak | yes |
| **S1-mFc** | 36 mg/L | yes | yes | <1 mg/L | yes | yes |
| **S1-His** | 15 mg/L | yes | yes | <1 mg/L | weak | no |
| **S1-S2-His** | 8 mg/L | yes | yes | <1 mg/L | no | no |

Max. production yields of SARS-CoV-2 spike protein/domains in insect cells (High Five) and mammalian cells (Expi293F). Proteins with His-tag produced in High Five cells and S1-hFc\* were additionally purified by SEC. \* with Furin site.

S1 as well as S1-S2 were more efficiently produced in insect cells compared to Expi293F cells. RBD-SD1 was produced well in both production systems. The binding of the produced spike domains/proteins to ACE2 was validated by ELISA and flow cytometry analysis on ACE2 positive cells (Table 1).

### Antibodies were selected by phage display

Antibodies were selected against SARS-CoV-2 spike S1 subunit in four panning rounds in microtiter plates. The following single clone screening was performed by antigen ELISA in 96 well MTPs, using soluble monoclonal scFv produced in *E. coli.* Subsequently, DNA encoding for the binders was sequenced and unique antibodies were recloned as scFv-Fc fusions.

In detail, three panning strategies were compared. In a first approach (STE70) the lambda (HAL9) and kappa (HAL10) libraries were combined and the antigen S1-hFc (with furin site, produced in High Five cells) was immobilized in PBS. Here, only seven unique antibodies were identified. In a second approach, the selection was performed separately for HAL10 (STE72) and HAL9 (STE73) using S1-hFc as antigen (with furin site, SEC purified, immobilized in carbonate buffer). Here, 90 unique antibodies were selected from HAL10 and 209 from HAL9. In a third approach (STE77 and STE78), S1-hFc produced in Expi293F cells was used (immobilized in carbonate buffer). Here, the panning resulted in only three unique antibodies that were not further analyzed in inhibition assays. An overview is given in Table 2.

**Table 2.** Antibody selection strategies using the human naive antibody gene libraries HAL9/10.

| Antibody selection campaign | library | target | panning rounds | binders/ screened clones | unique antibodies | cloned as scFv-Fc | inhibiting antibodies |
| --- | --- | --- | --- | --- | --- | --- | --- |
| STE70 | HAL9/10 | S1-hFc (Hi5) | 4 | 7/ 94 | 7 | 7 | 1 |
| STE72 | HAL10 (kappa) | S1-hFc (Hi5) | 4 | 397/ 752 | 90 | 44 | 8 |
| STE73 | HAL9 (lambda) | S1-hFc (Hi5) | 4 | 519/ 846 | 209 | 59 | 8 |
| STE77 | HAL10 (kappa) | S1-hFc (Expi) | 4 | 7/ 564 | 2 | n.a. | n.a. |
| STE78 | HAL9 (lambda) | S1-hFc (Expi) | 4 | 10/ 282 | 1 | n.a. | n.a. |

n.a. = not applicable

The antibody subfamily distribution was analyzed and compared to the subfamily distribution in the HAL9/10 library and *in vivo* (Fig. 1). The phage display selected antibodies mostly originated from the main gene families VH1 and VH3. Only few antibodies were found using VH4. In 96 of the 309 selected antibodies (31%), the V-gene VH3-23 was used. The V-gene distribution in the lambda light chains was similar to the distribution in the original library. Only antibodies comprising the V-gene VL6-57 were selected from the lambda library HAL10. In antibodies selected from the kappa library, VK2 and VK4 were underrepresented.

**Fig 1.**
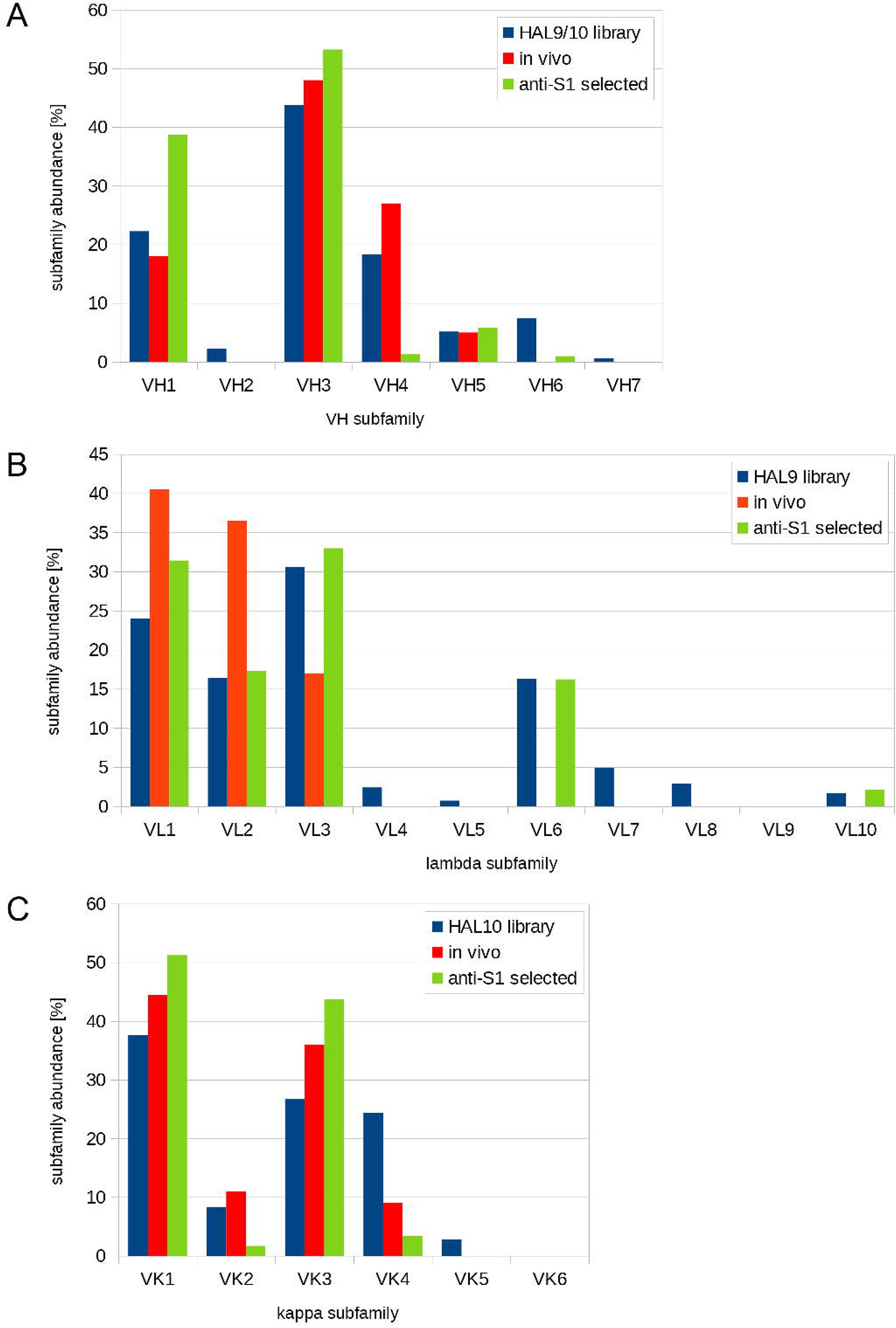
Use of V region genes in human anti-SARS-CoV-2 antibodies. Comparison of the distribution of V region gene subfamilies the in the universal HAL9/10 library ^50^, the *in vivo* distribution of subfamilies ^80^ and the distribution of antibodies against S1 selected from HAL9/10. (A) Abundance of VH, (B) Vκ and (C) Vλ.

### Anti-SARS-CoV-2 scFv-Fc were produced transiently in mammalian cells

In the interest of rapid throughput to quickly address the growing impact of the COVID-19 pandemic, only a selection of the unique antibodies was chosen for production as scFv-Fc and characterization. Antibodies with potential glycosylation sites in the CDRs, identified by *in silico* analysis, were excluded. A total of 109 scFv-Fc antibodies were produced in 5 mL culture scale, with yields ranging from of 20 to 440 mg/L.

### Antibodies inhibit the binding of spike to ACE2 positive cells in the scFv-Fc format

To further select potential therapeutic candidates, an inhibition assay was established using flow cytometry of ACE2-positive cells, measuring competition of S1-S2 trimer binding by scFv-Fc antibodies. The entire spike protein ectodomain was used for this inhibition assay for optimal representation of the viral binding. In a first screening, the 109 scFv-Fc were tested at 1500 nM (molar ratio antibody: S1-S2 30:1). 17 antibodies with inhibition better than 75% were selected for further analysis (Fig. 2A, Table 3 and Supplementary Data 3).

**Table 3.** Overview on inhibiting antibodies.

| Antibody name | VH | Germ-<br>inality<br>index<br>VH<br>[%] | VL | Germ-<br>inality<br>index<br>VL<br>[%] | EC50 ELISA [nM] scFv-Fc |  |  | EC50 ELISA [nM] IgG |  |  | flow cytometry scFv-Fc spike binding inhibition assay |  |  |  | scFv-Fc SARS-CoV-2 CPE based neutrali-<br>zation [%] |
| --- | --- | --- | --- | --- | --- | --- | --- | --- | --- | --- | --- | --- | --- | --- | --- |
|  |  |  |  |  | RBD | S1 | S1-S2 | RBD | S1 | S1-S2 | IC 50 [nM]<br>with 50 nM<br>spike | Molar ratio<br>antibody<br>arm: spike | IC 50 [nM]<br>with 10 nM<br>RBD | Molar ratio<br>antibody<br>arm: RBD |  |
| STE70-1E12 | VH1-2 | 96.7 | VL6-57 | 94.4 | 0.3 | 0.5 | 1.1 | 0.3 | 0.4 | 0.8 | 180 | 7.2 | 3.2 | 0.64 | 98 |
| STE72-1B6 | VH3-23 | 93.4 | VK1-12 | 95.5 | 0.5 | 0.7 | 1.4 | 1.1 | 1.6 | 2.4 | 240 | 9.6 | 4.8 | 0.96 | 90 |
| STE72-1G5 | VH1-69 | 98.9 | VK3-20 | 96.6 | 2.8 | 3.4 | 5.2 | n.a. |  |  | n.d. | n.d. | n.d. | n.d. | 77 |
| STE72-4C10 | VH3-30 | 97.8 | VK1D-39 | 92.1 | 0.5 | 1 | 2.4 | n.a. |  |  | 117 | 4.8 | 3.5 | 0.7 | 87 |
| STE72-4E12 | VH1-46 | 100 | VK3-15 | 98.9 | 1.5 | 3.3 | 3.7 | 1.6 | 1.1 | 1.7 | n.d. | n.d. | 3.0 | 0.6 | 99 |
| STE72-8A2 | VH1-18 | 100 | VK1D-33 | 97.8 | 0.5 | 0.7 | 0.8 | not binding as IgG |  |  | 35 | 1.4 | 1.5 | 0.3 | 97 |
| STE72-8A6 | VH1-18 | 100 | VK1-5 | 94.4 | 0.5 | 0.9 | 1.2 | not binding as IgG |  |  | 102 | 4.0 | 2.8 | 0.56 | 100 |
| STE72-8E1 | VH4-61 | 93.4 | VK1-5 | 93.3 | 0.4 | 0.8 | 0.6 | n.a. |  |  | 20 | 0.8 | 5.6 | 1.1 | 85 |
| STE72-2G4 | VH1-2 | 100 | VL2-8 | 94.3 | 0.2 | 0.3 | 0.3 | n.a. |  |  | 37 | 1.4 | 3.7 | 0.74 | 86 |
| STE73-2B2 | VH1-2 | 95.6 | VL6-57 | 92.2 | 0.2 | 0.3 | 0.3 | n.a. |  |  | 63 | 2.6 | 1.7 | 0.4 | 75 |
| STE73-2C2 | VH3-66 | 96.7 | VL6-57 | 92.2 | 3.1 | 5.7 | 7.8 | n.a. |  |  | 59 | 2.4 | 3.0 | 0.6 | 70 |
| STE73-2E9 | VH1-18 | 100 | VL1-36 | 96.6 | 0.2 | 0.2 | 0.2 | 0.2 | 0.2 | 0.2 | 20 | 0.8 | 3.4 | 0.68 | 90 |
| STE73-2G8 | VH3-66 | 92.3 | VL3-19 | 100 | 0.2 | 0.3 | 0.4 | 0.8 | 1.7 | 2.8 | 23 | 1.0 | 2.8 | 0.56 | 98 |
| STE73-6B10 | VH1-2 | 97.8 | VL2-11 | 94.3 | 5.5 | 4.9 | 20.2 | not binding as IgG |  |  | 612 | 24 | 73 | 14.6 | 90 |
| STE73-6C1 | VH3-30 | 98.9 | VL1-40 | 92.0 | 0.6 | 0.9 | 1.8 | 14.1 | 19.5 | 34 | 97 | 3.8 | 4.1 | 0.81 | 100 |
| STE73-6C8 | VH1-69 | 98.9 | VL6-57 | 93.3 | 1.1 | 1.9 | 5.4 | 2.9 | 4.3 | 4 | 332 | 13.2 | 5.4 | 1.08 | 100 |
| STE73-9G3 | VH3-23 | 97.8 | VL1-40 | 94.3 | 0.4 | 0.5 | 0.9 | 0.4 | 0.9 | 2.3 | 40 | 1.6 | 3.4 | 0.6 | 100 |

V-genes were determined by VBASE2 (vbase2.org) <sup>72</sup>. The EC50 were measured on 30 ng immobilized RBD-mFc, S1-mFc, S1-S2-His (trimer) by ELISA. The IC50 was measured by flow cytometry using 50 nM (in relation to monomer) S1-S2 trimer, respectively 10 nM RBD and ACE2 positive cells. The molar ration of antibody binding site: S1-S2 or RBD is given for 50% inhibition. n.a.: not applicable. CPE based neutralization assay was performed with 250 pfu/well SARS-CoV-2 and 1 µg/ml (~100 nM) (median neutralization %). EC50 were calculated with GraphPad Prism Version 6.1, fitting to a four-parameter logistic curve. IC50 values were calculated using Logistic5 fit of Origin.

**Fig 2.**
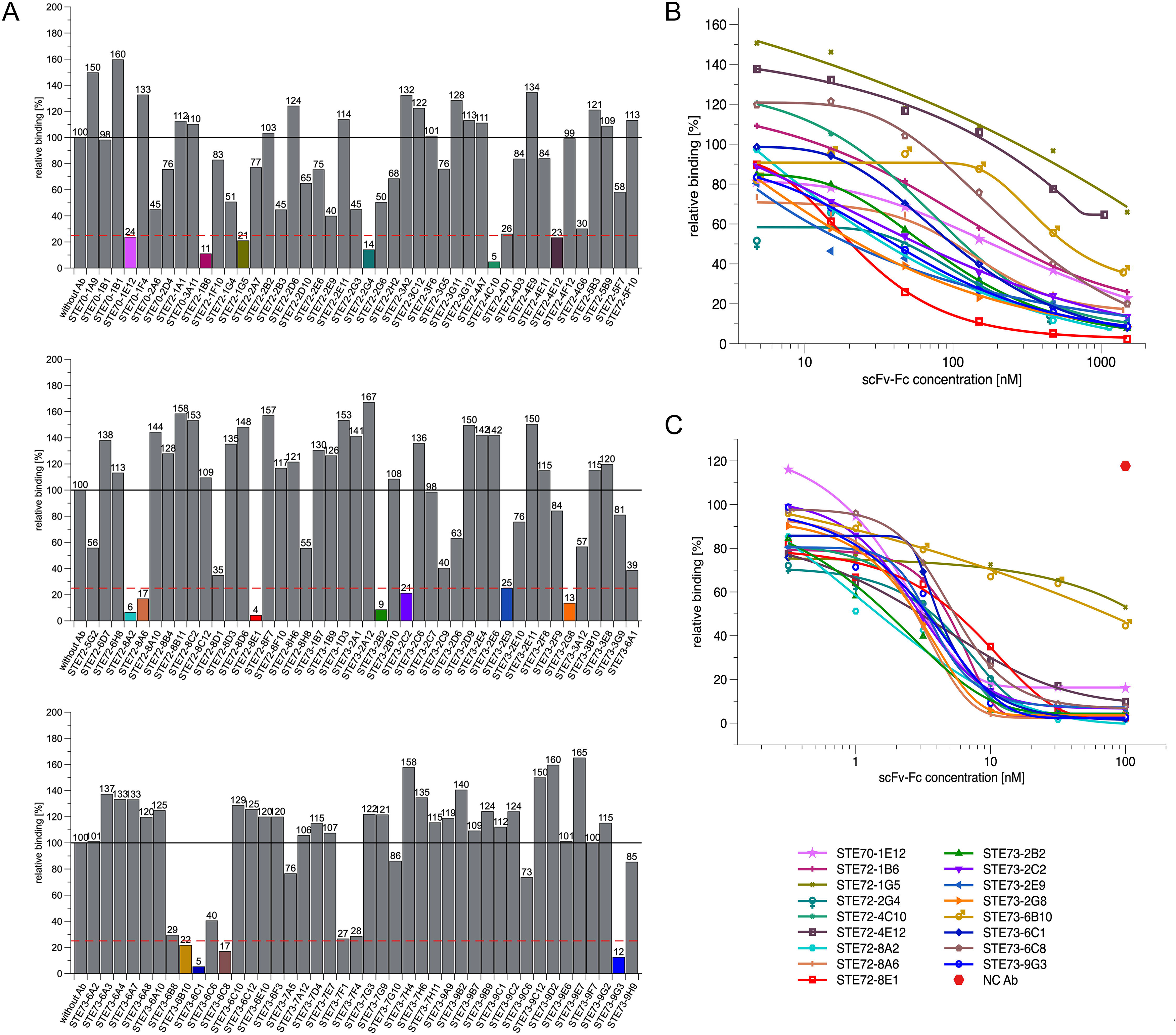
Inhibition of SARS-CoV-2 spike protein binding to cell (flow cytometry). (A) Inhibition prescreen of 109 scFv-Fc antibodies on ACE2 positive cells using 1500 nM antibody and 50 nM spike protein (30:1 ratio). The antibodies selected for detailed analysis are marked in colors. The antibodies STE73-9-G3 and STE73-7H10 (marked with *) are identical. (B) IC50 determination by flow cytometry using 50 nM S1-S2 trimer and 4.7 – 1500 nM scFv-Fc. (C) IC50 determination by flow cytometry using 10 nM RBD and 0.03-1000 nM scFv-Fc. Logistic5 fit of Origin was used to determine the IC50.

To further characterize these 17 antibodies, their inhibition of ACE2 binding was assessed at concentrations from 1500 nM to 4.7 nM (from 30:1 to ~1:10 Ab:antigen molar ratio) with the same flow cytometry assay (Fig. 2B and Table 3). Antibodies STE72-8E1 and STE73-2E9 showed 50% inhibition of ACE2 binding at a molar ratio of 0.8 antigen binding sites per spike monomer. For further validation of the direct RBD: ACE2 inhibition, we performed the same assay using a RBD-mFc construct (Fig. 2C). With the exception of two antibodies (STE72-1G5 and STE73-6B10) all antibodies showed high inhibition of binding with molar ratios of 0.3-0.6:1 for STE72-4E12, STE72-8A2, STE72-8A6, STE73-2B2, STE73-2G8 and STE73-9G3.

The inhibition of the 17 antibodies was further validated on human Calu-3 cells, which naturally express ACE2 ^9^ using RBD-mFc (Supplementary Data 4A) and S1-S2-His (Supplementary Data 4B) showing a stronger inhibition on Calu-3 compared to the transiently overexpressing ACE2 positive Expi293F cells. The Expi293F system allowed an improved estimation of inhibition potency when using the complete S1-S2 spike protein, because the S1-S2 was directly labeled with a fluorophore and the signals were not amplified in comparison to RBD with a murine Fc and a fluorophore labeled secondary antibody. Further, ACE2-expressing Expi293F cells present a much higher amount of ACE2-receptor on their surface compared to Calu-3, due to the CMV-mediated expression (data not shown). Taken together these data show that all 17 inhibiting antibodies selected against S1 directly interfered with RBD-ACE2 binding.

### Determination of EC50 of the inhibiting antibodies to RBD, S1 and S1-S2

The EC50 of the inhibiting scFv-Fc on RBD, S1 (without furin site) and S1-S2 spike was measured by ELISA. All inhibiting antibodies bound the isolated RBD (Fig. 3), identifying it as their target on the viral surface. Most of the inhibiting antibodies showed a half-maximal binding in the subnanomolar range for RBD. While STE72-2G4 showed sub-nanomolar EC50 values for RBD and S1, it was discarded due to noticeable cross-reactivity to mFc. The EC50 on the S1-S2 spike trimer was reduced for most of the antibodies, in comparison to the isolated RBD or S1.

**Fig 3.**
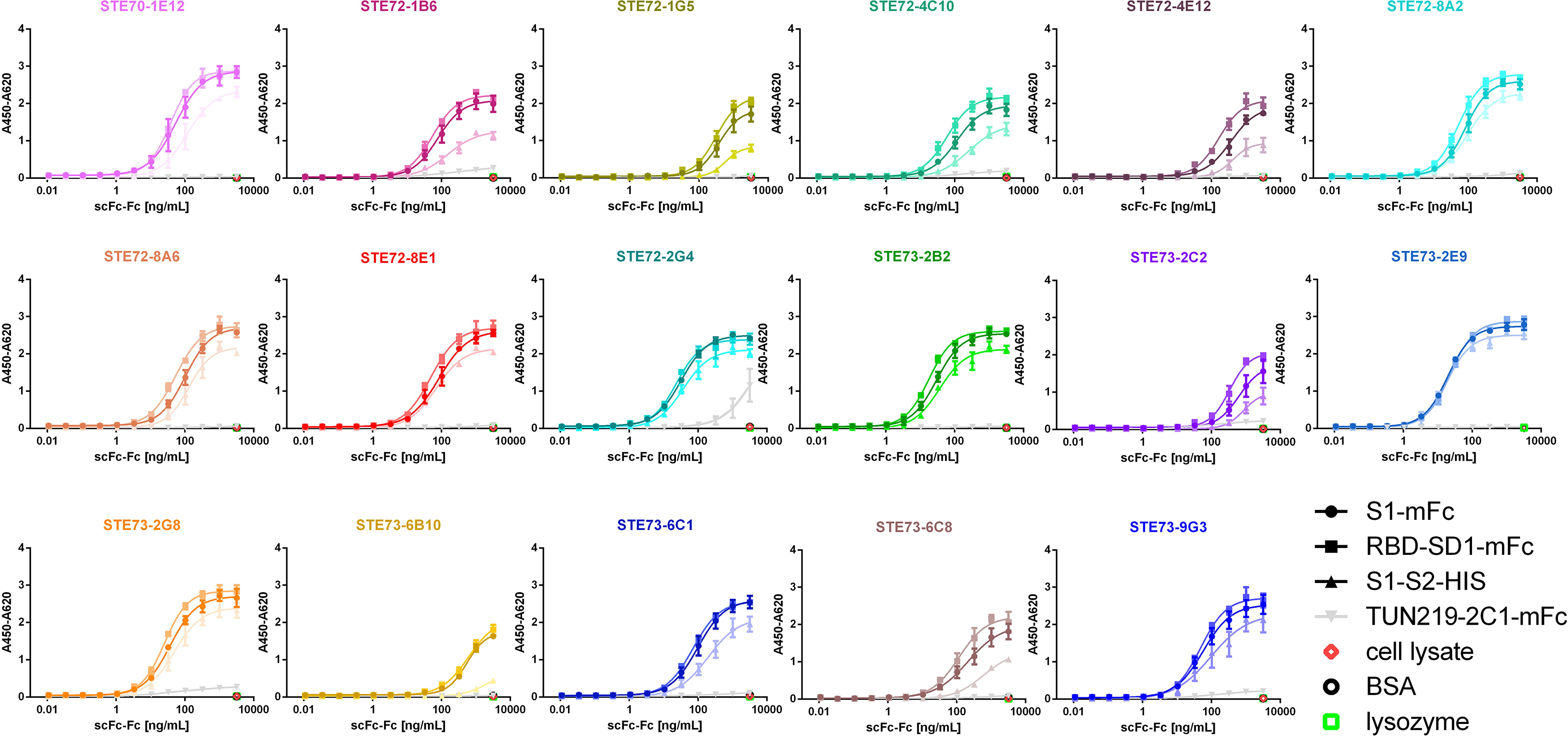
Determinination of EC50 on RBD. Binding in titration ELISA of the 17 best inhibiting scFv-Fc on RBD (fusion protein with murine Fc part), S1 (fusion protein with murine Fc part) or S1-S2 (fusion protein with His tag). Sequence SARS-CoV-2 (Gene bank QHD43416). An unrelated antibody with murine Fc part (TUN219-2C1), human HEK293 cell lysate, BSA or lysozyme were used as controls. EC50 were calculated with GraphPad Prism Version 6.1, fitting to a four-parameter logistic curve.

### ScFv-Fc combinations show synergistic effects in inhibition assays

Combinations of best-inhibiting scFv-Fc were tested in the flow cytometry inhibition assay using 1500 nM antibody and 50 nM S1-S2 spike (Supplementary Data 5). Some of the combinations showed an increase of inhibition compared to the same amount of individual antibodies.

### Anti-RBD scFv-Fc neutralize active SARS-CoV-2

All 17 inhibiting scFv-Fc were screened in a cytopathic effect (CPE)-based neutralization assay using 250 pfu/well SARS-CoV-2 Münster/FI110320/1/2020 and 1 μg/mL (~10 nM) scFv-Fc (Fig. 4A, Table 3). VeroE6 cells showed pronounced CPE characterised by rounding and detachment clearly visible in phase contrast microscopy upon SARS-CoV-2 infection within 4 days, while uninfected cells maintained an undisturbed confluent monolayer. Virus inoculum pre-incubated with anti-RBD antibodies led to decreased CPE in varying degrees quantified by automated image analysis for cell confluence. All 17 antibodies showed neutralization in this assay. Fig. 4B shows examples for strong (STE73-6C8) and weak (STE73-2C2) neutralizing antibodies and controls.

**Fig 4.**
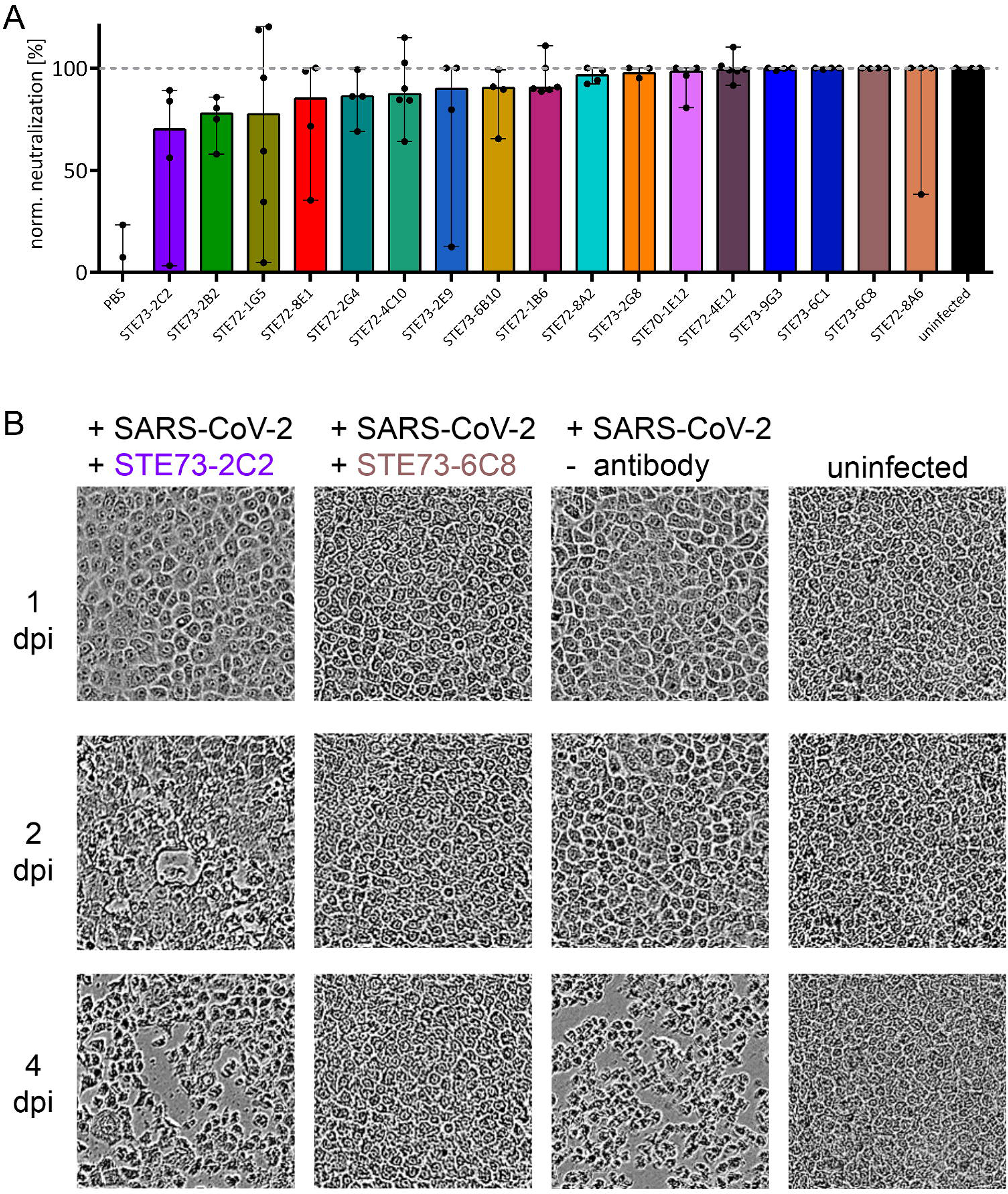
SARS-CoV-2 neutralization in the scFv-Fc format. Neutralization analysis using 250 pfu of SARS-CoV-2 in a CPE based neutralization assay. A) Cell monolayer occupancy at 4 days post infection in absence of neutralizing antibodies was compared to uninfected control cells and median values were normalized as 0 and 100% occupancy, respectively. Histograms indicate medians of normalized monolayer occupancy in a neutralization assay using 1 μg/mL (~10 nM) antibody for each of the 17 tested antibodies. Black dots indicate monolayer occupancy in individual assays (4-6 measurements per sample). B) Representative phase contrast microscopy pictures of uninfected cells, cells infected in absence of antibodies, in the presence of a poorly neutralizing scFv-Fc (STE73-2C2) or of a highly neutralizing scFv-Fc (STE73-6C8).

### Binding, inhibition and SARS-CoV-2 neutralization in the IgG format

Eleven scFv-Fc showing a neutralization efficacy of 90% according to the CPE-based neutralization assay were converted into the IgG format. First, their binding was analyzed by titration ELISA on RBD, S1 and S1-S2 (Supplementary Data 6, Table 3). Three antibodies lost binding after conversion to IgG (STE72-8A2, STE72-8A6 and STE73-6B10, data not shown), others showed reduced binding of different degrees, while three antibodies retained their binding (STE70-1E12, STE72-4E12 and STE73-2E9). In the next step, the antibodies were tested in the cell-based inhibition assay using RBD (Figure 5A) or S1-S2 (Figure 5B). Here, the inhibition was confirmed for STE73-2E9, −9G3, −2G8. STE73-1B6 showed inhibition of RBD comparable to the latter antibodies, but its activity was almost absent on S1-S2, thus it was not further considered.

**Fig 5.**
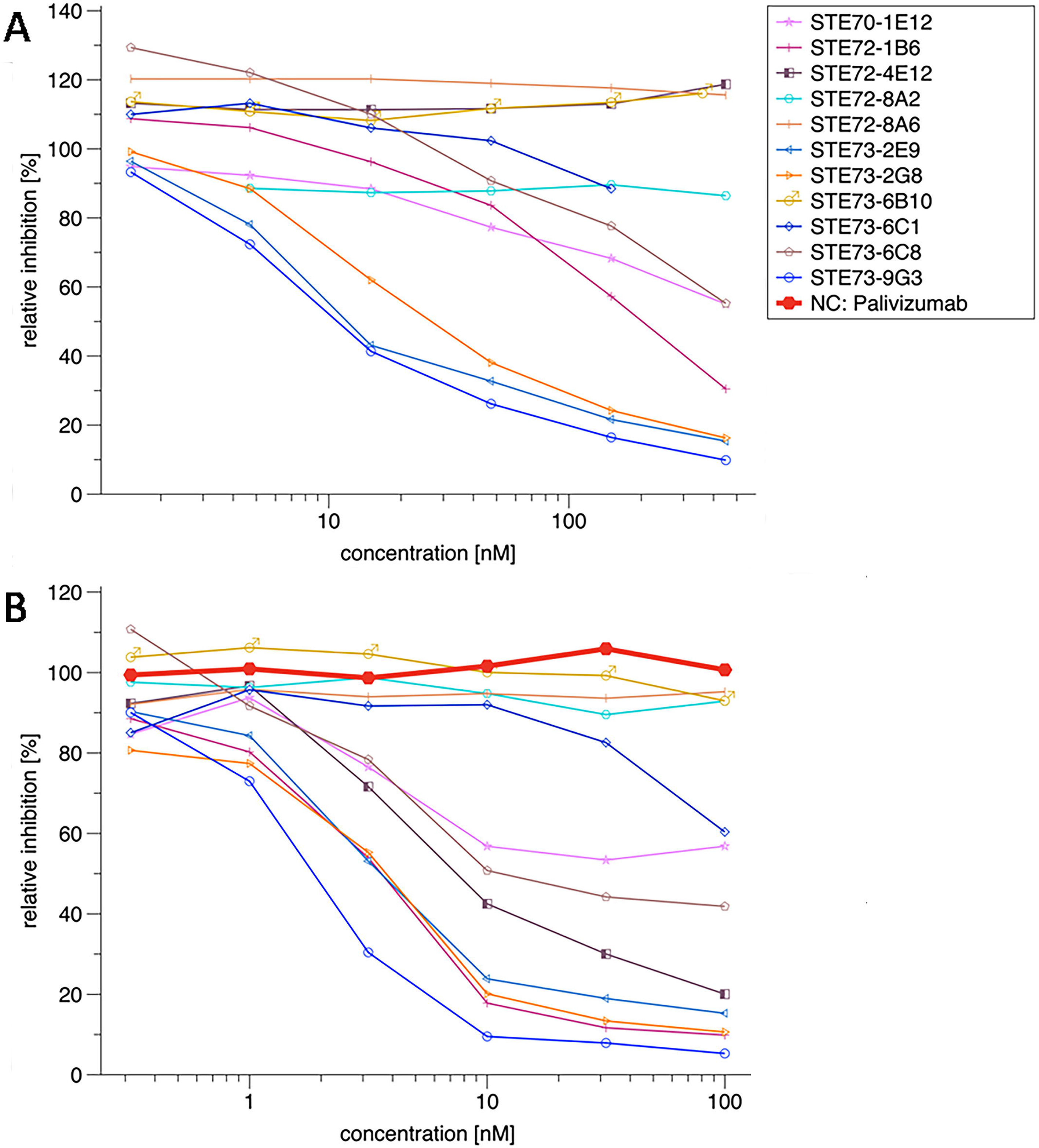
Inhibition of RBD-ACE2 interaction by IgG. (A) IC50 determination by flow cytometry using 50 nM S1-S2-His and 0.5-500 nM IgG. (B) IC50 determination by flow cytometry using 10 nM RBD-mFC and 0.1-100 nM IgG. Palivizumab was used as negative control. Logistic5 fit of Origin was used to determine the IC50.

### Inhibiting IgGs are binding at the RBD-ACE2 interface

The efficiently inhibiting IgGs STE73-2E9, STE73-2G8 and STE73-9G3 were analyzed for their binding to various S1 subunit variants harbouring a panel of recently reported mutations in RBD region and the D614G mutation. Three assays were employed: 1, ELISA (Figure 6A), 2, surface plasmon resonance (SPR) and 3, bScreen protein array with S1 proteins from different sources. All three antibodies lost binding to RBD mutations in the region aa483-486 directly at the RBD-ACE2 interface, showing that mutations in that region affect their epitope on the antigen. There were only minor differences between different approaches, e.g. at positions aa439 and aa476 for STE73-2E9. We then used this information to guide and validate computational docking simulations followed by atomistic molecular dynamics simulations according to protocols developed and well established in our group ^35^, obtaining three-dimensional atomic models of the antibody-RBD interaction for these three antibodies (Figure 6B). The binding models of the three antibodies to the spike ectodomain are shown in Supplementary Data 7.

**Fig 6.**
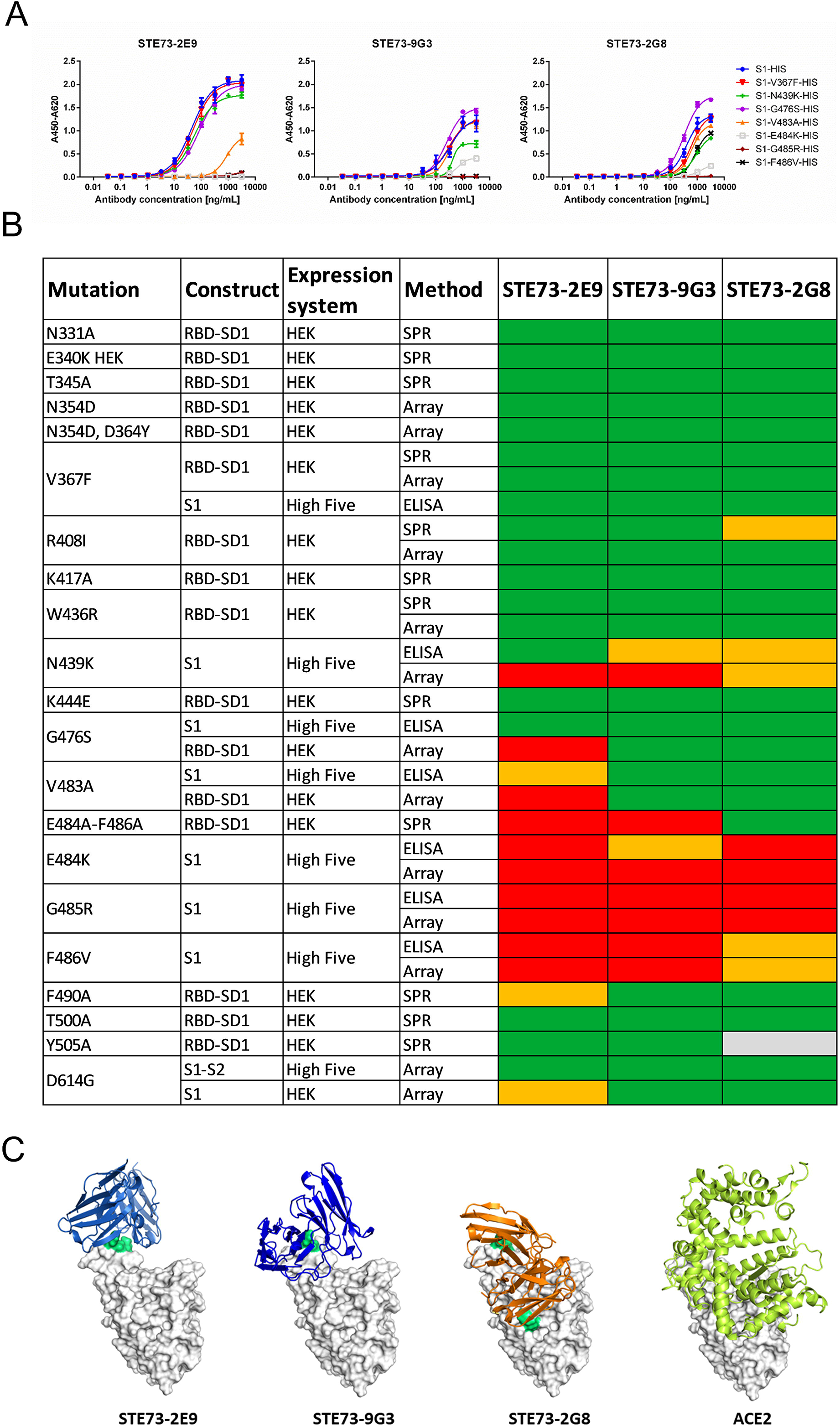
Binding to RBD mutants, epitopes and structure models. (A) ELISA using STE73-2E9, −9G3 and −2G8 on S1-His with different RBD mutations. (B) Overview of the binding of STE73-2E9, −9G3 and −2G8 to different RBD mutations analyzed by ELISA, SPR and protein array. Sequence SARS-CoV-2 (Gene bank QHD43416). (C) The three antibodies STE73-2E9, −9G3 and −2G8 are binding to the ACE-RBD interface (docking models based on epitope data from binding to RBD mutations). Experimentally validated computational models of the variable regions of the antibodies (coloured cartoons) binding to the RBD (white surface, same orientation in all images) are shown. The cartoon representation of ACE2 is also shown for comparison.

### STE73-2E9 neutralize SARS-CoV-2 in the IgG format

As a final step, STE73-2E9, −9G3, −2G8 antibodies were analyzed in a plaque assay using the patient isolate SARS-CoV-2 (Münster/FI110320/1/2020) to determine their neutralization potency. STE73-2E9 showed an IC50 of 0.43 nM in this assay. Unfortunately, STE73-9G3 and −2G8 did not show high neutralization as IgG (Figure 7A).

**Fig 7.**
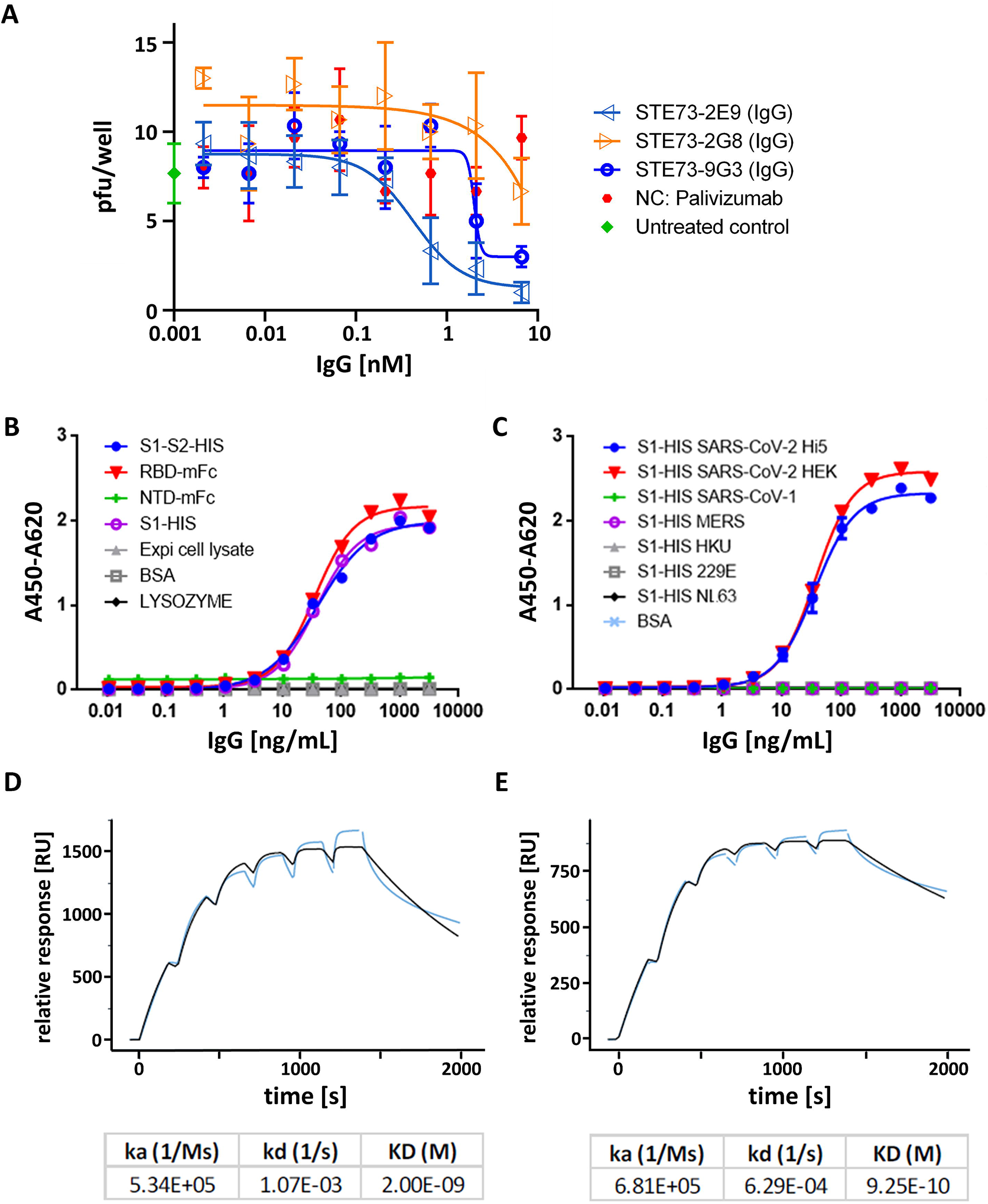
Characterization of the neutralizing antibody STE73-2E9 in IgG format. (A) Neutralization of 20-30 pfu SARS-CoV-2 by STE73-2E9, −9G3 and −2G8. Palivizumab was used as isotype control. (B) Titration ELISA on the indicated antigens. (C) Cross-reactivity to other Coronavirus spike proteins analzyed by ELISA. S1-HIS SARS-CoV-2 Hi5 was produced in house. S1-HIS SARS-CoV-2 HEK and all other coronaviruses S1 domain proteins were obtained commercially. (D and E) Kinetic parameters determination through single cycle kinetic titration SPR of STE73-2E9 IgG on HEK cell produced RBD-SD1 and S1-S2, respectively (concentrations: 200, 100, 50, 25, 12.5, 6.25 nM).

### Analysis of STE73-2E9 crossreactivity with other coronoviruses

The neutralizing antibody STE73-2E9 was further characterized by titration ELISA on SARS-CoV-2 spike recombinant constructs (Figure 7B) and S1 subunits from different coronaviruses (Figure 7C) showing that STE73-2E9 is binding specifically SARS-CoV-2 Spike protein in the RBD region. The specific binding to S1/RBD of SARS-CoV-2 was further confirmed by the bScreen protein array binding analysis (data not shown).

### STE73-2E9 binds with nM affinity to RBD

The affinity of STE73-2E9 was determined by Surface Plasmon Resonance as 2×10^−9^ M for RBD-SD1 (Figure 7D) and 9.25^−10^ M for the complete spike protein (Figure 7E).

### Aggregation behaviour of STE73-2E9

The aggregation behaviour of biologicals is a key factor for therapeutic development. STE73-2E9 shows now relevant aggregation under normal conditions (pH7.4, RT in PBS), heat stress conditions (pH7.4, 45°C, 24h in PBS) and pH stress (pH3, 24h, RT), implicating that it has benign general physicochemical properties that are a prerequisite for the development into a passive vaccine (supplementary data 8).

## Discussion

For 130 years, antibodies in animal sera or convalescent human plasma were successfully used for the treatment of infectious diseases, starting with the work of Emil von Behring und Shibasaburo Kitasato against diphtheria ^35^. However, the efficacy of human plasma derived from convalescent donors depends on the viral pathogen. In case of Ebola, the survival upon treatment with convalescent human plasma was not significantly improved over the control group ^36^. On the other hand, reduced mortality and safety was shown for convalescent plasma transfer in case of influenza A H1N1 in 2009 ^37,38^. This approach was also used against emerging coronaviruses. While the outcomes were not significantly improved in a very limited number of MERS patients ^39^, the treatment was successful for SARS ^40,41^. This approach was also used for COVID-19 with promising results ^42^. The mode of action of these polyclonal antibody preparations may vary, including virus neutralization, Fcγ receptor binding mediated phagocytosis or antibody dependent cellular cytotoxicity as well as complement activation ^43–45^. In any serum therapy, the composition and efficacy of convalescent plasma is expected to differ from donor to donor, as well as batch to batch, and sera must be carefully controlled for viral contaminations (e.g. HIV, Hepatitis viruses) and neutralization potency. A convalescent patient can provide 400-800 mL plasma, with 250-300 mL of plasma typically needed per treatment. With two rounds of treatment per patient, this is a grave limitation, since one donor can only provide material for 1-2 patients ^42,43^. Human or humanized monoclonal antibodies are a powerful alternative to polyclonal antibodies derived from convalescent plasma. Following this approach, the humanized antibody Palivizumab was approved in 2009 for treatment and prevention of Respiratory Syncytial Virus (RSV) infections ^46^. Other antibodies against viral diseases successfully tested in clinical studies are mAb114 and REGN-EB3 against Ebola disease ^24^.

Phage display derived antibodies are typically well established medications: twelve such antibodies are approved by EMA/FDA at the time of writing, a significant increase compared to the six such antibodies approved in 2016 ^47^. In this work, we used phage display to isolate monoclonal human antibodies capable of neutralizing SARS-CoV-2 from a universal, naive antibody gene library that was generated from healthy donors before the emergence of the SARS-CoV-2 virus. This allowed selection of human antibodies against this virus without the necessity to obtain material from COVID-19 infected individuals. While most antibodies against SARS-CoV-2 were obtained from convalescent patients with few exceptions ^20,21,48,49^ our approach demonstrates that human antibodies with functional properties matching those of the antibodies isolated from convalescent patients can be generated without the necessity to wait for material from COVID-19 infected individuals. Therefore, this strategy offers a very fast additional opportunity to respond to future pandemics.

As the human receptor of SARS-CoV and SARS-CoV-2 is ACE2 ^3^, we focused on antibodies which directly block the interaction of the spike protein with this receptor and antibodies preventing ACE2 binding were shown to potently neutralize the closely related SARS-CoV virus ^16^. 309 unique fully human monoclonal antibodies were generated using different panning strategies. The S1 subunit produced in insect cells was better suited for antibody selection than the S1 subunit produced in mammalian cells. The V-gene distribution of the selected anti-Spike antibodies is largely in accordance with the V-gene subfamily distribution shown by Kügler et al ^50^ for antibodies selected against 121 other antigens from HAL9/10. Only the VH1 subfamily was over-represented and VH4 and Vkappa4 subfamilies were rarely selected despite their presence in the HAL libraries. The most frequently used V-gene was VH3-30. Interestingly, an increased use of this V-gene in anti-SARS-CoV-2 antibodies was also described by Robbiani *et al.* 2020 ^51^ for anti-RBD B-cells selected from COVID-19 patients. By contrast, the second most selected V-gene was VH3-53, which was selected in our approach only once. Robbiani *et al.* also described an overrepresented use of VL6-57, as found in our antibodies as well. However, it has to be noted that VL6 is also overrepresented in our naïve library compared to its *in vivo* occurrence.

From the initial 309 scFv, 109 were recloned in the scFv-Fc IgG-like bivalent format. Their ability to inhibit binding of fluorescently labelled S1-S2 trimer to ACE2 expressing cells was assessed by flow cytometry. The half-maximal inhibition of the best inhibiting 17 scFv-Fc was measured both with the spike trimer and isolated RBD. Significantly, some of the antibodies showed half-maximal inhibition at a ratio around 1:1 - in certain cases even better - when calculated per individual binding site (antigen binding site:spike monomer/RBD). A similar molar ratio of 1:1 was demonstrated by Miethe *et al*. 2014 for inhibition of botulinum toxin A ^52^. In the trimeric spike protein, the RBD can be in an “up” (open) or “down” (close) position. The “down” conformation can not bind to ACE2, in contrast to the less stable “up” conformation ^33^. The RBDs can be in different conformations on the same spike trimer, which offers a possible explanation for the observed effective antibody to spike molar ratios lower than 1:1. This is in accordance with the cryo-EM images recorded by Walls et al. ^4^, where they could find half of the recorded trimers with one RBD in the open conformation. We observed that molar ratios for half maximal inhibition were lower for RBD compared to spike protein. For some antibodies, approximately 0.5 antigen binding sites were needed to achieve a 50% inhibition. The fact that the antibodies are more efficient at inhibiting RBD binding to ACE2 rather than S1-S2 trimer binding can be explained with the higher affinity of the antibodies for the isolated RBD compared to the trimeric spike, which in turn points to the presence of partially or completely inaccessible epitopes on the trimer, an occurrence seen in other viruses. This is similar to what was reported by Pinto *et al.* ^17^ who also showed a lower affinity of the antibody S309 for spike compared to RBD.

Inhibition of ACE2 binding was stronger on the human lung cells Calu-3, which better represent the *in vivo* situation than transiently ACE2 overexpressing cells. Nevertheless, we did the titration assays on ACE2 overexpressing Expi293F cells because these seemed to allow a better quantitative discrimination of inhibiting potency.

Antibody combinations can have a synergistic effect as previously described for toxins and viruses ^32,53–55^. This approach may also avoid formation of viral escape mutants. Here, the best combinations showed a significantly improved inhibition efficacy, at least when using an excess of antibodies (Ab:Agmolar ratio 30:1).

All of the 17 scFv-Fc were tested in neutralization assays using a SARS-CoV-2 strain isolated from a patient and all antibodies showed a degree of neutralization in this assay. While this study did not aim to define the lowest effective concentration of individual antibodies in limiting dilution conditions, all tested antibodies showed a clear and measurable effect at a relatively low concentration. Therefore, our approach provided a rapid selection of antiviral antibodies.

In a next step, we converted eleven antibodies with the best neutralization efficacy according to the cytopathic assay into the IgG format. It was completely unexpected that most antibodies lost efficacy in the inhibition assay after conversion from scFv-Fc to IgG including antibodies like STE70-1E12 without loss of affinity according to the titration ELISA. These results are in contrast to former results where none ^54^ or only a low percentage ^55,56^ of antibodies lost efficacy after conversion from scFv-Fc to IgG. Nevertheless, three antibodies showed a good inhibition in the cell-based assay and did not bind to the region of aa483-486 known for RBD mutations from publications ^19,57^ and from GISAID database (www.gisaid.org). Experimentally validated computational docking shows that these antibodies still at least partially occupy the ACE2 binding site on the RBD, thus likely achieving direct inhibition of virus-ACE2 interaction. The binding sites of antibody BD368-2 ^18^, B38 ^58^ and REGN10933 ^59^ also overlap with the RBD ACE2 binding interface. The neutralizing antibody STE73-2E9 was specific for SARS-CoV-2 and we conclude that this antibody is a suitable candidate for the development of passive immunotherapy for the treatment of COVID-19. It could be used therapeutically to prevent individuals from being hospitalized in intensive care units (ICUs), but also prophylactically, to protect health care workers or risk groups that do not respond to vaccination. Before clinical application, the risk of antibody-dependent enhancement of disease (ADE) has to be considered for COVID-19. In contrast to antibodies against Ebola where ADCC is important for protection ^22^, antibodies directed against the spike protein of SARS-CoV-2 may lead to ADE ^60–62^. SARS cause an acute lung injury which is also driven by immune dysregulation and inflammation caused by anti-spike antibodies ^63^. While, Quinlan *et al*. ^64^ described that animals immunized with RBD SARS-CoV-2 did not mediate ADE and suggested for vaccines the use of RBD, some of the monoclonal antibodies we analyzed in this study lead to an increased binding of the spike protein to ACE2 positive cells. A possible explanation could be multimerization of the spike by antibody ‘cross-linking’ in this assay or the stabilization of an infection-promoting conformation by the antibodies. These aspects need to be carefully considered in any development of therapeutic antibodies, we suggest to focus on RBD and/or the use of silenced Fc parts with deleted Fcγ and C1q binding ^65–67^ for safety reasons.

In conclusion, we report the successful isolation and characterization of a fully human, recombinant anti-spike neutralizing monoclonal antibody from naïve phage display libraries. Our approach demonstrates how neutralizing antibodies can be efficiently selected in a rapid time frame and without the need of convalescent patient material. Furthermore, the strategy we used efficiently targeted the spike:ACE interface allowing the selection of directly blocking antibodies.

## Methods

### Design of expression vectors

Production in Expi293F cells was performed using pCSE2.5-His-XP, pCSE2.6-hFc-XP or pCSE2.6-mFc-XP ^68^ where the respective single chain variable fragment of the antibodies or antigens were inserted by *Nco*I/*Not*I (NEB Biolabs) digestion. Antigen production in High Five insect cells was performed using *Nco*I/*Not*I compatible variants of the OpiE2 plasmid ^69^ containing an N-terminal signal peptide of the mouse Ig heavy chain, the respective antigen and C-terminal either 6xHis-tag, hFc or mFc. Single point mutations in S1-HIS constructs were inserted through site-directed mutagenesis using overlapping primers according to Zheng et al. ^70^ with slight modifications: S7 Fusion polymerase (Mobidiag, Espoo, Finland) with the provided GC buffer and 3%DMSO was used for the amplification reaction.

### Production of antigens in insect cells

Different domains or subunits of the Spike protein (GenBank: MN908947), S1 subunit mutants and the extracellular domain of ACE2 receptor (GenBank NM_021804.3) were Baculovirus-free produced in High Five cells (Thermo Fisher Scientific) by transient transfection as previously described in Bleckmann *et al.* ^34^. Briefly, High Five cells were cultivated at 27°C, 110 rpm in ExCell405 media (Sigma) and kept at a cell density between 0.3 – 5.5 ×10^6^ cells/mL. For transfection cells were centrifuged and resuspended in fresh media to a density of 4×10^6^ cells/mL and transfected with 4 μg plasmid/mL and 16 μg/mL of PEI 40 kDa (Polysciences). 4 h up to 24 h after transfection cells were fed with 75% of the transfection volume. At 48 h after transfection cell culture medium was doubled. Cell supernatant was harvested five days after transfection in a two step centrifugation (4 min at 180xg and 20 min at above 3500xg) and 0.2 μm filtered for purification.

### Production of antigens and scFv-Fc in mammalian cells

Antibodies, different domains or subunits of the Spike protein and the extracellular domain of ACE2 were produced in Expi293F cells (Thermo Fisher Scientific). Expi293F cells were cultured at 37°C, 110 rpm and 5% CO2 in Gibco FreeStyle F17 expression media (Thermo Fisher Scientific) supplemented with 8 mM Glutamine and 0.1% Pluronic F68 (PAN Biotech). At the day of transfection cell density was between 1.5 - 2×10^6^ cells/mL and viability at least above 90%. For formation of DNA:PEI complexes 1 μg DNA/mL transfection volume and 5 μg of 40 kDa PEI (Polysciences) were first diluted separately in 5% transfection volume in supplemented F17 media. DNA and PEI was then mixed and incubated ~25 min at RT before addition to the cells. 48 h later the culture volume was doubled by feeding HyClone SFM4Transfx-293 media (GE Healthcare) supplemented with 8 mM Glutamine. Additionally, HyClone Boost 6 supplement (GE Healthcare) was added with 10% of the end volume. One week after transfection supernatant was harvested by 15 min centrifugation at 1500xg.

### Protein purification

Protein purification was performed depending on the production scale in either 24 well filter plate with 0.5 mL resin (10 mL scale) or 1 mL column on Äkta go (Cytiva), Äkta Pure (Cytiva) or Profina System (BIO-RAD). MabSelect SuRe or HiTrap Fibro PrismA (Cytiva) was used as resins for Protein A purification. For His-tag purification of Expi293F supernatant HisTrap FF Crude column (Cytiva) and for His-tag purification of insect cell supernatant HisTrap excel column (Cytiva) was used. All purifications were performed according to the manufactures manual. Indicated antigens were further purified by size exclusion chromatography by a 16/600 Superdex 200 kDa pg (Cytiva). All antigens, antibodies and scFv-Fc were run on Superdex 200 Increase 10/300GL (Cytiva) on Äkta or HPLC (Techlab) on an AdvanceBio SEC 300Å 2.7 μm, 7.8×300 mm (Agilent) for quality control.

### Validation of spike protein binding to ACE2

ACE2 binding to the produced antigens was confirmed in ELISA (enzyme linked immunosorbent assay) and on cells in flow cytometry. For ELISA, 200 ng ACE2-mFc per well was immobilized on a Costar High binding 96 well plate (Corning, Costar) at 4°C over night. Next, the wells were blocked with 350 μL 2% MBPST (2% (w/v) milk powder in PBS; 0.05% Tween20) for 1 h at RT and then washed 3 times with H_2_O and 0.05% Tween20 (BioTek Instruments, EL405). Afterwards, the respective antigen was added at the indicated concentrations and incubated 1 h at RT prior to another 3 times washing step. Finally, the antigen was detected using mouse-anti-polyHis conjugated with horseradish peroxidase (HRP) (1:20000, A7058, Sigma) for His-tagged antigens, goat-anti-mIgG(Fc) conjugated with HRP (1:42000, A0168, Sigma) for mFc tagged antigen versions or goat-anti-hIgG(Fc) conjugated with HRP (1:70000, A0170, Sigma) if hFc-tagged antigens had to be detected. Bound antigens were visualized with tetramethylbenzidine (TMB) substrate (20 parts TMB solution A (30 mM Potassium citrate; 1 % (w/v) Citric acid (pH 4.1)) and 1 part TMB solution B (10 mM TMB; 10% (v/v) Acetone; 90% (v/v) Ethanol; 80 mM H_2_O_2_ (30%)) were mixed). After addition of 1 N H_2_SO_4_ to stop the reaction, absorbance at 450 nm with a 620 nm reference wavelength was measured in an ELISA plate reader (BioTek Instruments, Epoch).

To verify the ACE2-antigen interaction on living cells, Expi293F cells were transfected according to the protocol above using pCSE2.5-ACE2_fl_-His and 5% eGFP plasmid. Two days after transfection, purified S1-S2-His, S1-His or RBD-His were labelled using Monolith NT^TM^ His-Tag Labeling Kit RED-tris-NTA (Nanotemper) according to the manufacturer’s protocol. Fc-tagged ligand versions were labelled indirectly by using goat-anti-mFc-APC (Dianova) or mouse anti-hFcγ-APC (Biolegend) antibody. 100, 50 and 25 nM of antigen were incubated with 5×10^5^ ACE2-expressing or non-transfected Expi293F cells (negative control) 50 min on ice. After two washing steps, fluorescence was measured in MACSQuant Analyzer (Miltenyi Biotec.).

### Antibody selection using phage display

The antibody selection was performed as described previously with modifications ^71^. In brief, for panning procedure, the antigen was immobilized on a High binding 96 well plate (Corning, Costar). 5 μg of S1-hFc (produced in High Five cells) was diluted in carbonate puffer (50 mM NaHCO_3_/Na_2_CO_3_, pH 9.6) and coated onto the wells at 4°C overnight. Next, the wells were blocked with 350 μL 2% MBPST (2% (w/v) milk powder in PBS; 0.05% Tween20) for 1 h at RT and then washed 3 times with PBST (PBS; 0.05% Tween20). Before adding the libraries to the coated wells, the libraries (5×10^10^ phage particles) were preincubated with 5 μg of an unrelated scFv-Fc and 2% MPBST on blocked wells for 1 h at RT, to deprive libraries of human Fc fragment binders. The libraries were transferred to the antigen-coated wells, incubated for 2 h at RT and washed 10 times. Bound phage were eluted with 150 μL trypsin (10 μg/mL) at 37°C, 30 minutes and used for the next panning round. The eluted phage solution was transferred to a 96 deep well plate (Greiner Bio-One, Frickenhausen, Germany) and incubated with 150 μL *E. coli* TG1 (OD_600_ = 0.5) firstly for 30 min at 37°C, then 30 min at 37°C and 650 rpm to infect the phage particles. 1 mL 2xYT-GA (1.6% (w/v) Tryptone; 1 % (w/v) Yeast extract; 0.5% (w/v) NaCl (pH 7.0), 100 mM D-Glucose, 100 μg/mL ampicillin) was added and incubated for 1 h at 37°C and 650 rpm, followed by addition of 1×10^10^ cfu M13KO7 helper phage. Subsequently, the infected bacteria were incubated for 30 min at 37°C followed by 30 min at 37°C and 650 rpm before centrifugation for 10 min at 3220xg. The supernatant was discarded and the pellet resuspended in fresh 2xYT-AK (1.6% (w/v) Tryptone; 1 % (w/v) Yeast extract; 0.5% (w/v) NaCl (pH 7.0), 100 μg/mL ampicillin, 50 μg/mL kanamycin). The antibody phage were amplified overnight at 30°C and 650 rpm and used for the next panning round. In total four panning rounds were performed. In each round, the stringency of the washing procedure was increased (20x in panning round 2, 30x in panning round 3, 40x in panning round 4) and the amount of antigen was reduced (2.5 μg in panning round 2, 1.5 μg in panning round 3 and 1 μg in panning round 4). After the fourth as well as third panning round single clones containing plates were used to select monoclonal antibody clones for the screening ELISA.

### Screening of monoclonal recombinant binders using E. coli scFv supernatant

Soluble antibody fragments (scFv) were produced in 96-well polypropylene MTPs (U96 PP, Greiner Bio-One) as described before ^55,71^. Briefly, 150 μL 2xYT-GA was inoculated with the bacteria bearing scFv expressing phagemids. MTPs were incubated overnight at 37°C and 800 rpm in a MTP shaker (Thermoshaker PST-60HL-4, Lab4You, Berlin, Germany). A volume of 180 μL 2xYT-GA in a PP-MTP well was inoculated with 20 μL of the overnight culture and grown at 37°C and 800 rpm for 90 minutes (approx. OD _600_ of 0.5). Bacteria were harvested by centrifugation for 10 min at 3220xg and the supernatant was discarded. To induce expression of the antibody genes, the pellets were resuspended in 200 μL 2xYT supplemented with 100 μg/mL ampicillin and 50 μM isopropyl-beta D thiogalacto pyranoside (IPTG) and incubated at 30°C and 800 rpm overnight. Bacteria were pelleted by centrifugation for 20 min at 3220xg and 4°C.

For the ELISA, 100 ng of antigen was coated on 96 well microtiter plates (High binding, Greiner) in PBS (pH 7.4) overnight at 4°C. After coating, the wells were blocked with 2% MPBST for 1 h at RT, followed by three washing steps with H_2_O and 0.05% Tween20. Supernatants containing secreted monoclonal scFv were mixed with 2% MPBST (1:2) and incubated onto the antigen coated plates for 1 h at 37°C followed by three H_2_O and 0.05% Tween20 washing cycles. Bound scFv were detected using murine mAb 9E10 which recognizes the C-terminal c-myc tag (1:50 diluted in 2% MPBST) and a goat anti-mouse serum conjugated with horseradish peroxidase (HRP) (A0168, Sigma) (1:42000 dilution in 2% MPBST). Bound antibodies were visualized with tetramethylbenzidine (TMB) substrate (20 parts TMB solution A (30 mM Potassium citrate; 1% (w/v) Citric acid (pH 4.1)) and 1 part TMB solution B (10 mM TMB; 10% (v/v) Acetone; 90% (v/v) Ethanol; 80 mM H_2_O_2_ (30%)) were mixed). After stopping the reaction by addition of 1 N H_2_SO_4_, absorbance at 450 nm with a 620 nm reference was measured in an ELISA plate reader (Epoch, BioTek). Monoclonal binders were sequenced and analyzed using VBASE2 (www.vbase2.org) ^72^ and possible glycosylation positions in the CDRS were analyzed according to Lu et al ^73^.

### Inhibition of S1-S2 binding to ACE2 expressing cells using MacsQuant

The inhibition tests in cytometer on EXPI293F cells were performed based on the protocol for *validation of spike protein binding to ACE2* (see above) but only binding to S1-S2-His and RBD-mFc antigen (High Five cell produced) was analyzed. The assay was done in two setups. In the first setup 50 nM antigen was incubated with min. 1 μM of different scFv-Fc and the ACE2 expressing cells. The resulting median antigen fluorescence of GFP positive living single cells was measured. For comparison of the different scFv-Fc first the median fluorescence background of cells without antigen was subtracted, second it was normalized to the antigen signal where no antibody was applied. All scFv-Fc showing an inhibition in this setup were further analyzed by titration (max. 1500 nM-4.7 nM) on S1-S2-His (High Five cell produced), respectively on RBD-mFc (max. 100 nM-0.03 nM). The IC50 was calculated using the equation f(x)=Amin+(Amax-Amin)/(1+(x0/x)^h)^s and parameters from Origin. In addition, pairwise combinations (max. 750 nM of each scFv-Fc) of the different inhibiting scFv-Fc were tested.

### Dose dependent binding of the antibodies (scFv-Fc or IgG format) in titration ELISA

ELISA were essentially performed as described above in “S*creening of monoclonal recombinant binders using E.coli scFv supernatant”*. For titration ELISA the purified scFv-hFc were titrated from 3.18 μg/mL-0.001 ng/mL on 30ng/well of the following antigens: S1-S2-His (High Five cell produced), RBD-mFc (High Five cell produced), S1-mFc (High Five cell produced) and TUN219-2C1-mFc (as control for unspecific Fc binding). In addition, all scFv-hFc were also tested only at the highest concentration (3.18 μg/mL) for unspecific cross-reactivity on Expi293F cell lysate (10^4^cells/well), BSA (1% w/v) and lysozyme. ScFv-hFc or IgG were detected using goat-anti-hIgG(Fc)-HRP (1:70000, A0170, Sigma). Titration assays were performed using 384 well microtiter plates (Costar) using Precision XS microplate sample processor (BioTek), EL406 washer dispenser (BioTek) and BioStack Microplate stacker (BioTek). EC50 were calculated with by GraphPad Prism Version 6.1, fitting to a four-parameter logistic curve. The binding of antibodies to S1 subunit His-tagged constructs containing mutations in the RBD region and the cross-reactivity to Spike proteins of other coronaviruses was tested as described above. S1-HIS proteins from SARS-CoV-2 (expressed in HEK cells), SARS-CoV-1, MERS, HCoV HKU1, HCoV NL63, HCoV 229E were acquired commercially (Sino Biologicals products 40591-V08H, 40150-V08B1, 40069-V08H, 40021-V08H, 40601-V08H, 40600-V08H).

### Antibody structures and computational docking studies

The antibody structures were modelled according to the canonical structure method using the RosettaAntibody program ^74^ as previously described ^75^ and docked to the experimental structure of SARS-CoV-2 S protein receptor binding domain (RBD, PDBid: 6M17) ^6^.

Docking was performed using the RosettaDock 3.12 software ^76^ as previously described ^77^. Briefly, each antibody was manually placed with the CDR loops facing the RBD region containing the residues identified by the peptide mapping experiment. The two partners were moved away from each other by 25Å and then brought together by the computational docking algorithm, obtaining thousands of computationally generated complexes (typically 15,000). The antibody/RBD complexes were structurally clustered and then selected according to the scoring function (an estimate of energetically favourable solutions) and agreement with the peptide mapping data. Selected complexes were further optimized by a docking refinement step and molecular dynamics simulations.

### SARS-CoV-2 neutralization in cell culture

VeroE6 cells (ATCC CRL-1586) were seeded at a density of 6*10^4^/well onto cell culture 96-well plates (Nunc, Cat.#167008). Two days later, cells reached 100% confluence. For neutralization, antibodies (1 μg/ml final concentration) were mixed with the virus inoculum (250 pfu/well), using strain SARS-CoV-2/Münster/FI110320/1/2020 (kind gift of Stephan Ludwig, University of Münster, Germany), in 100 μl full VeroE6 culture medium (DMEM, 10% FCS, 2 mM glutamine, penicillin, streptomycin) in technical quadruplicates or sixfold replicates and incubated for 1 hour at 37°C. Then, cells were overlaid with the antibody/virus mix and phase contrast images were taken automatically using a Sartorius IncuCyte S3 (10x objective, two hours image intervals, 4 images per well) housed in a HeraCell 150i incubator (37°C, 100% humidity, 5% CO2). Image data was quantified with the IncuCyte S3 GUI tools measuring the decrease of confluence concomitant with the cytopathic effect of the virus in relation to uninfected controls and controls without antibody and analyzed with GraphPad Prism 8. Given is the median of the inhibition.

For titration, antibodies were diluted in 1/√10 steps and mixed with a fixed inoculum of SARS-CoV-2 (100-150 pfu) in a total volume of 500 μl of Vero E6 medium. After one hour incubation at 37°C, cells were infected with the antibody/virus mix, incubated for one hour and then overlaid with Vero E6 medium containing 1.5% methyl-cellulose. Three days postinfection, wells were imaged using a Sartorius IncuCyte S3 (4x objective, whole-well scan) and plaques were counted from these images.

### Cloning and production of IgG

For IgG production, selected antibodies were converted into the human IgG1 format by subcloning of VH in the vector pCSEH1c (heavy chain) and VL in the vector pCSL3l/pCSL3k (light chain lambda/kappa) ^78^, adapted for Golden Gate Assembly procedure with Esp3I restriction enzyme (New England Biolabs). EXPI293F (Thermo Fisher Scientific) cells were transfected with 12.5 μg of both vectors in parallel in a 1:1 ratio. For production, the transfected EXPI293F cells were cultured in chemically defined medium F17 (Thermo Fisher Scientific) supplemented with 0.1% pluronic F68 (PAN-Biotech,) and 7.5 mM L-glutamine (Merck) for seven days. A subsequent protein A purification was performed as described above.

### Affinity determination by surface plasmon resonance (SPR)

The antibodies binding properties were analyzed at 25 °C on a Biacore ™ 8K instrument (GE Healthcare) using 10mM HEPES pH 7.4, 150mM NaCl, 3mM EDTA and 0.005% Tween-20 as running buffer. SARS-CoV2 RBDs, wild-type and different mutants, or full S-protein, were immobilized on the surface of a CM5 chip through standard amine coupling. Increasing concentration of antibodies (6.25-12.5-25-50-100nM) were injected using a Single-cycle kinetics setting and analyte responses were corrected for unspecific binding and buffer responses. Curve fitting and data analysis were performed with Biacore™ Insight Evaluation Software.

### Analysis of binding to RBD mutants by protein array

2 nL of the proteins were printed as quadruplicates onto PDITC coated bScreen slides with a pitch of 700μm using a SciFlexArrayer (Scienion AG) in non-contact mode. The following proteins were spotted on the array: S1-S2, SARS-CoV-2, High5 (this work); S1, SARS-CoV-2, HEK293 (Sino Biological 40591-V08H); S1, SARS-CoV-2, Baculovirus (Sino Biological 40591-V08B1); S1-RBD, SARS-CoV-2, HEK293 (Sino Biological 40592-V08H); S1-humFc, SARS-CoV-2, HEK293 (Sino Biological 40591-V02H); S1S2, SARS-CoV-2, Baculovirus (Sino Biological 40589-V08B1); S2, SARS-CoV-2, Baculovirus (Sino Biological 40590-V08B); S1-RBD mFc, SARS-CoV-2, HEK293 (Sino Biological 40592-V05H); S1-RBD32, SARS-CoV-2, High5 (this work); S1-RBD25, SARS-CoV-2, High5 (this work); S1-RBD22, SARS-CoV-2, High5 (this work); S2, SARS-CoV-2, High5 (this work); S1, SARS-CoV-2, High5 (this work); S1 mFc, SARS-CoV-2, High5 (this work); S1-RBD32 mFc, SARS-CoV-2, High5 (this work); S1(E484K), SARS-CoV-2, High5 (this work); S1(F486V), SARS-CoV-2, High5 (this work); S1(N438K), SARS-CoV-2, High5 (this work); S1(G458R), SARS-CoV-2, High5 (this work); S1S2(D614G), SARS-CoV-2, High5 (this work); S1-RBD(V367F), SARS-CoV-2, HEK293 (Sino Biological 40592-V08H1); S1-RBD(V483A), SARS-CoV-2, HEK293 (Sino Biological 40592-V08H5); S1-RBD(R408I), SARS-CoV-2, HEK293 (Acro Biosystems SPD-S52H8); S1-RBD(G476S), SARS-CoV-2, HEK293 (Acro Biosystems SPD-C52H4); S1-RBD(N354D, D364Y), SARS-CoV-2, HEK293 (Acro Biosystems SPD-S52H3); S1-RBD(N354D), SARS-CoV-2, HEK293 (Acro Biosystems SPD-S52H5); S1-RBD(W436R), SARS-CoV-2, HEK293 (Acro Biosystems SPD-S52H7); S1-RBD, SARS-CoV-2, Baculovirus (Sino Biological 40592-V08B); S1(D614G), SARS-CoV-2, HEK293 (Sino Biological 40591-V08H3);; S1-S2, SARS, HEK293 (Acro Biosystems SPN-S52H5); S1, SARS, HEK293 (Acro Biosystems SIN-S52H5); S1-RBD, SARS, HEK293 (Acro Biosystems SPD-S52H6); S1-RBD, MERS, Baculovirus (Sino Biological 40071-V08B1); S1, MERS, HEK293 (Sino Biological 40069-V08H); N, MERS, Baculovirus (Sino Biological 40068-V08B); S1, HCoV-229E, HEK293 (Acro Biosystems SIN-V52H4); S1, HCoV-NL63, HEK293 (Acro Biosystems SIN-V52H3); S1, HCoV-HKU1, HEK293 (Sino Biological 40602-V08H); Streptavidin Cy5 (Thermo Fisher 434316); bBSA, (Thermo Fisher 29130), BSA (Carl Roth GmbH 8076.4). Protein concentration was adjusted to 200μg/mL, with the following exceptions: S1-RBD(E484K) 50 μg/mL, S1-RBD(F486V) 100 μg/mL, S1-RBD(N439K) 138 μg/mL, S1-RBD(G485R) 91 μg/ml. A bscreen was used for label-free measurement of antibody binding to the protein arrays. PBS BSA (1mg/ml) was used as sample as well as washing buffer. The flow-rate was set to 3μL/s. The measurement was performed in 3 steps - 1st step blocking solution (50% PBS BSA 1/mg/mL, 50% Superblock (Thermo Scientific - Cat No: 37515)); 2nd step neutralizing SARS-CoV-2 antibody 8μg/mL; 3rd step goat anti human Alexa 546 antibody 5μg/mL (Invitrogen - Cat No: A-21089). Each step consisted of 150s baselining, 333s association, 300s dissociation. The label-free signals of the association phase of the anti human step were used for data generation. This was done by subtracting the signal mean value from 30s to 20s before the association phase from the signal mean value from 20s to 30s after the association phase.

### Antibody structures and computational docking studies

The antibody structures were modelled according to the canonical structure method using the RosettaAntibody program ^74^ as previously described ^75^ and docked to the experimental structure of SARS-CoV-2 S protein receptor binding domain (RBD, PDBid: 6M17) ^6^.

Docking was performed using the RosettaDock 3.12 software ^76^ as previously described ^77^. Briefly, each antibody was manually placed with the CDR loops facing the RBD region containing the residues identified by the peptide mapping experiment. The two partners were moved away from each other by 25Å and then brought together by the computational docking algorithm, obtaining thousands of computationally generated complexes (typically 15,000). The antibody/RBD complexes were structurally clustered and then selected according to the scoring function (an estimate of energetically favourable solutions) and agreement with the peptide mapping data. Selected complexes were further optimized by a docking refinement step and molecular dynamics simulations.

The MD simulations were performed using GROMACS ^79^ with standard MD protocol: antibody/antigen complexes were centered in a triclinic box, 0.2 nm from the edge, filled with SPCE water model and 0.15M Na+Cl− ions using the AMBER99SB-ILDN protein force field; energy minimization followed. Temperature (298K) and pressure (1 Bar) equilibration steps of 100ps each were performed. 500ns MD simulations was run with the above-mentioned force field for each protein complexes. MD trajectory files were analyzed after removal of Periodic Boundary Conditions. The overall stability of each simulated complex was verified by root mean square deviation, radius of gyration and visual analysis according to standard procedures.

## Acknowledgements

We kindly acknowledge the support of the European Union for the ATAC (“antibody therapy against corona”, Horizon2020 number 101003650) consortium and the MWK Niedersachsen (14-76103-184 CORONA-2/20). L. Varani gratefully acknowledges support from SNF and Lions Club Monteceneri. Work was also supported by the Deutsche Forschungsgemeinschaft (DFG), grant KFO342 to L. Brunotte and S. Ludwig. We would like to highlight the passion and motivation of the complete team working on this topic in this special time. We are deeply grateful to Adelheid Langner, Andrea Walzog, Bettina Sandner, Cornelia Oltmann and Wolfgang Grassl for constant help and support.

## Author contributions

F.B., Gü.R., S.D., L.V., L.Č-Š., M.S., M.H. conceptualized the study. F.B., D.M., N.L., S.S., U.R., L.S., P.A.H., R.B., M.R., K-T.S., K.D.R.R., P.R., K.E., Y.K., D.S., M.P., S.Z.E., J.W., N.K., T.H., M.B., M.G., S.D.K., Gü.R., M.S. performed and designed experiments. F.B., D.M. N.L., S.S., U.R., L.S., P.K., Gü.R., L.V., L.Ĉ-Ŝ., M.S., M.H. analyzed data. S.L., L.B. provided material. S.D., L.V., L.Ĉ-Ŝ., M.H. conceived the funding. P.K., E.V.W., Gi.R., A.K., V.F., S.D., M.S. advised on experimental design and data analysis. F.B., S.D., Gü.R., L.V., L.Ĉ-Ŝ., M.S., M.H. wrote the manuscript.

