## Supplementary Data for "SARS-CoV-2 neutralizing human recombinant antibodies selected from pre-pandemic healthy donors binding at RBD-ACE2 interface"

**Supplementary Data 1** Schematic overview of the expressed variants of the Spike SARS-CoV-2 protein.

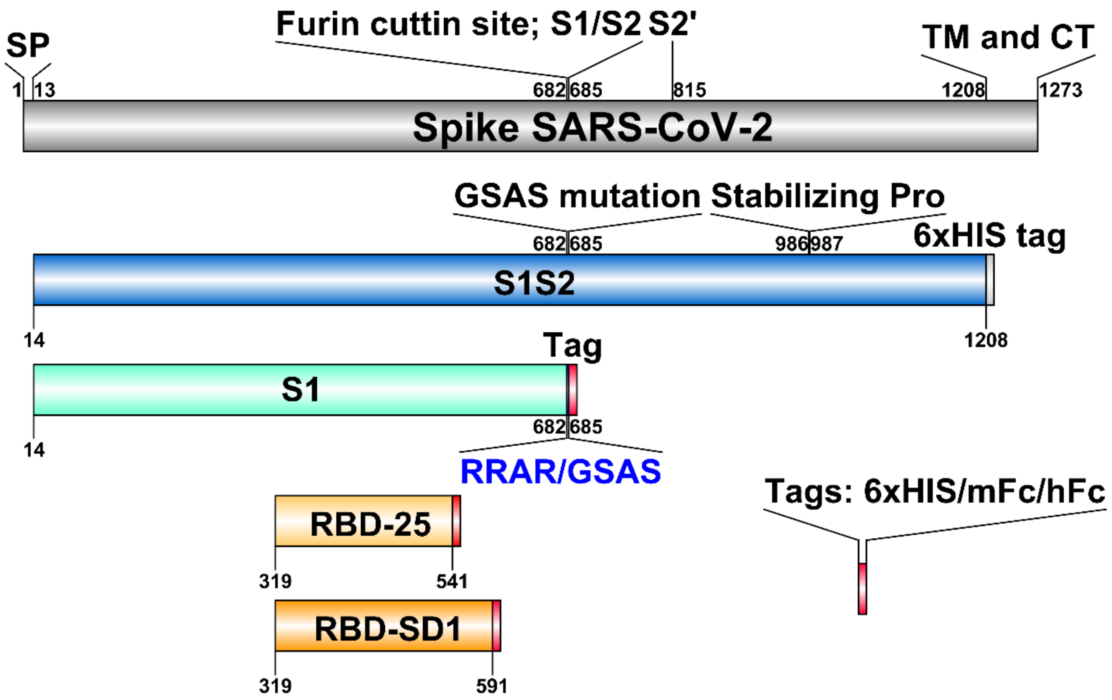

**Supplementary Data 2** SEC data of antigen produced in insect High Five cells (A-C) and mammalian Expi293F cells (D-E). The table (F) indicates the most likely conformation of the protein due to the retention volume of the peaks with the corresponding area in percentage. (*Furin site is present)

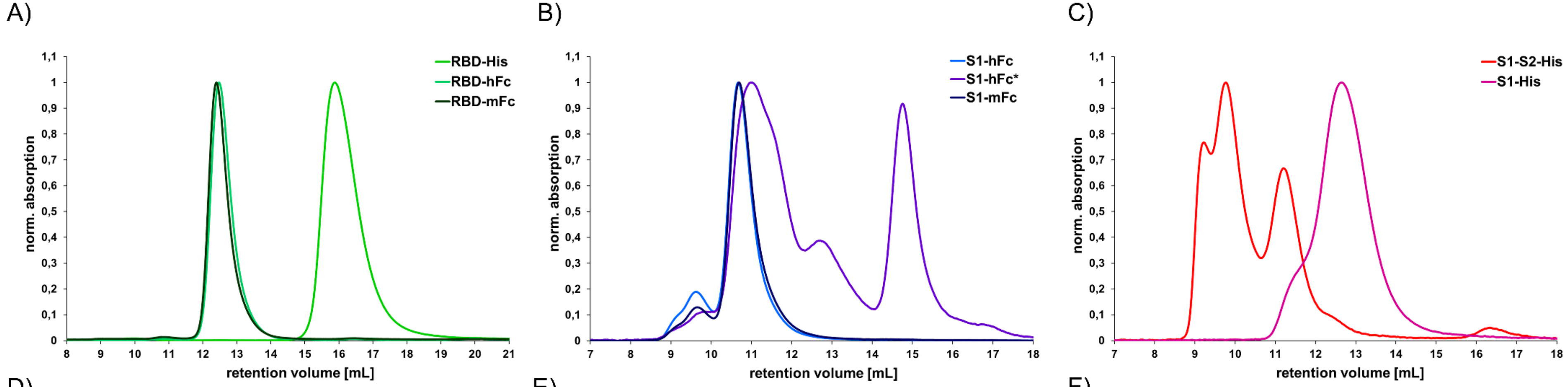


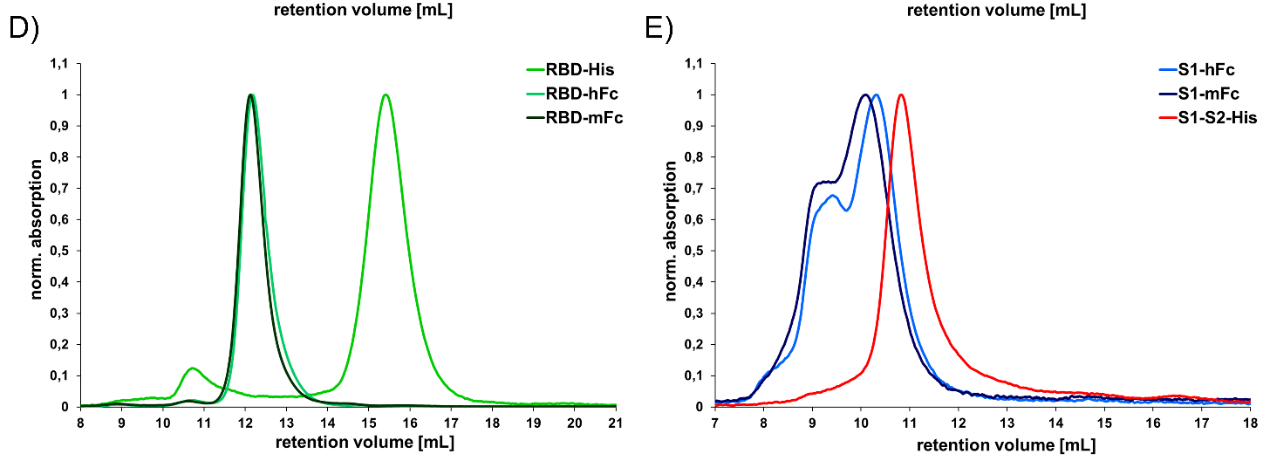


F)

| Antigen | Main Peak | High Five cells Vol [mL] | High Five cells Area [%] | Expi293F cells Vol [mL] | Expi293F cells Area [%] |
|---|---|---|---|---|---|
| RBD-His | Monomer | 15.9 | 99 | 15.4 | 93 |
| RBD-hFc | Dimer | 12.5 | 100 | 12.2 | 98 |
| RBD-mFc | Dimer | 12.4 | 99 | 12.1 | 99 |
| S1-His | Monomer | 12.7 | 100 | nd | nd |
| S1-hFc | Aggregate? | 9.6 | 16 | 9.3 | 31 |
| | Dimer | 10.7 | 84 | 10.3 | 68 |
| S1-hFc* | Dimer | 10.9 | 58 | nd | nd |
| | S1-hFc-hFc | 12.8 | 9 | nd | nd |
| | hFc Dimer | 14.8 | 33 | nd | nd |
| S1-mFc | Aggregate? | 9.7 | 11 | 9.1 | 27 |
| | Dimer | 10.7 | 89 | 10.1 | 73 |
| S1-S2-His | Aggregate? | 9.2 | 23 | - | - |
| | Trimer | 9.8 | 50 | - | - |
| | Monomer | 11.2 | 27 | 10.8 | 100 |

**Supplementary Data 3** Flow cytometry inhibition analysis A) Gateing strategy B) Analysis of spike S1-S2 trimer (50 nM in relation to monomer) binding to living cells expressing ACE2 blocked by 1500 nM antibodies and C) of RBD-mFc (10 nM in relation to monomer) binding to living cells expressing ACE2 blocked by 100 nM antibody. The antibodies STE72-1G5, STE72-4C10, STE72-8E1 and STE73-6C8 were used as example. The background control are transfected ACE2 cells and the reference are ACE2 positive cells incubated with labeled spike protein.

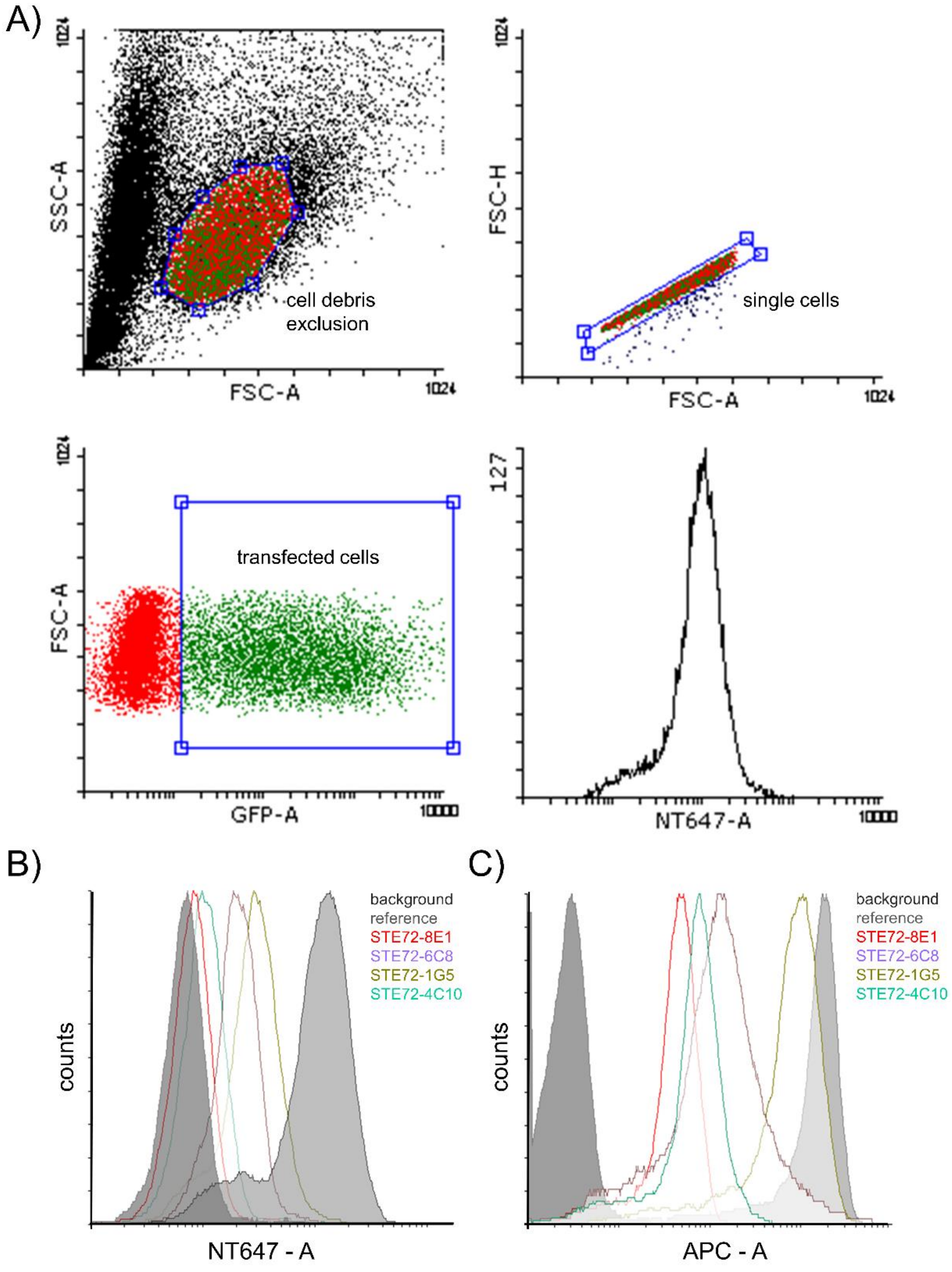

**Supplementary Data 4** Flow cytometry inhibition analysis A) Comparison of the inhibition of spike protein and RBD by flow cytometry on ACE2 positive Expi293F cells using 1000 nM scFv-Fc and 50 nM spike protein, respectively RBD (20:1 ratio). B) Inhibition of RBD binding in comparison of Expi293F transiently expressing ACE2 and Calu-3 cells. C) Inhibition of spike S1-S2 binding in comparison of Expi293F transiently expressing ACE2 and Calu-3. As negative control, the antibody SH1351-C1 was used.

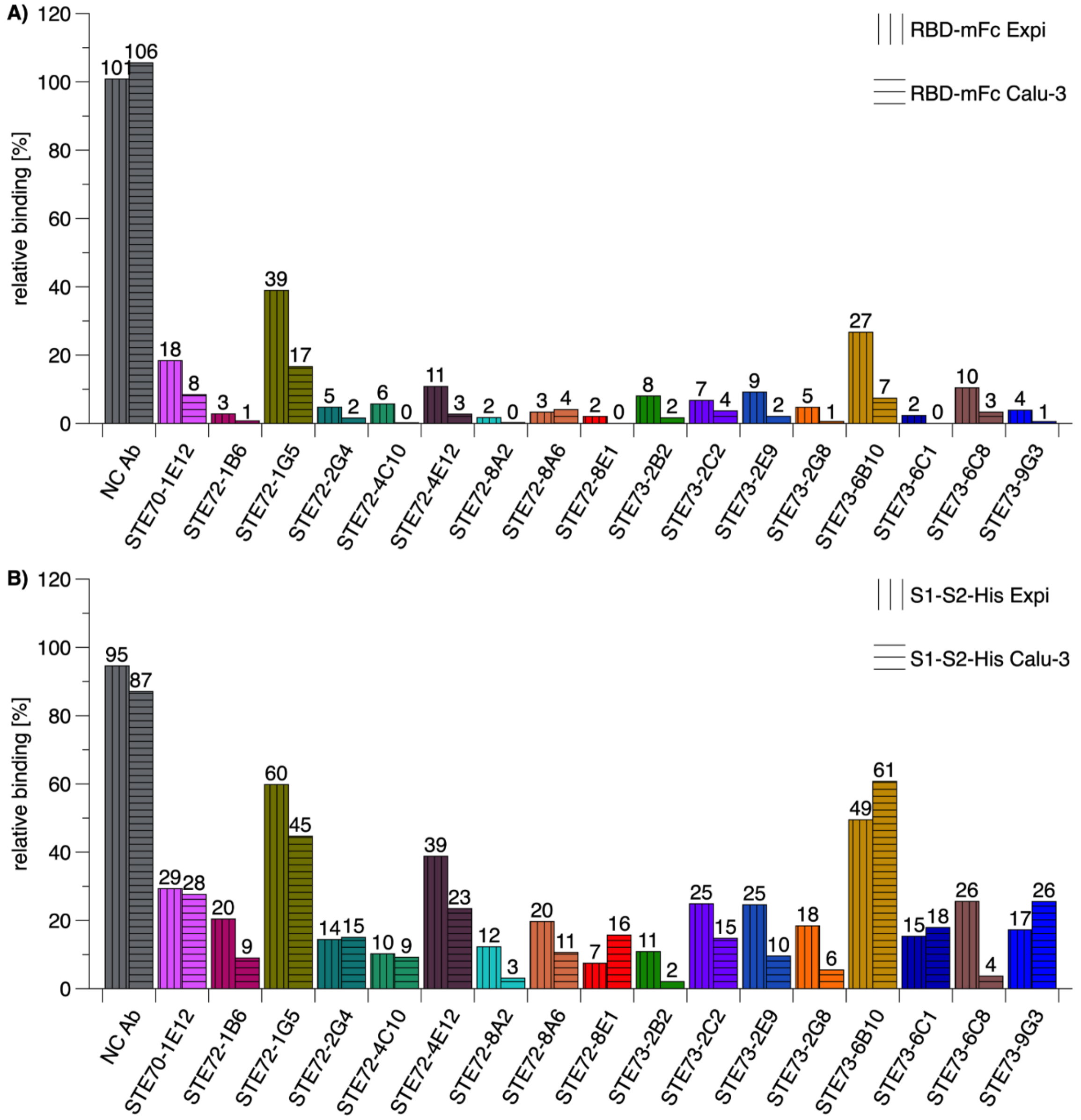

**Supplementary Data 5** Inhibition of SARS-CoV-2 spike protein binding to cells by antibody combinations. Flow cytometry analysis to determine the inhibition efficacy of antibody combinations on ACE2 positive cells and 50 nM S1-S2 and 1500 nM as a single antibody, respectively 750 nM of each antibody in a combination.

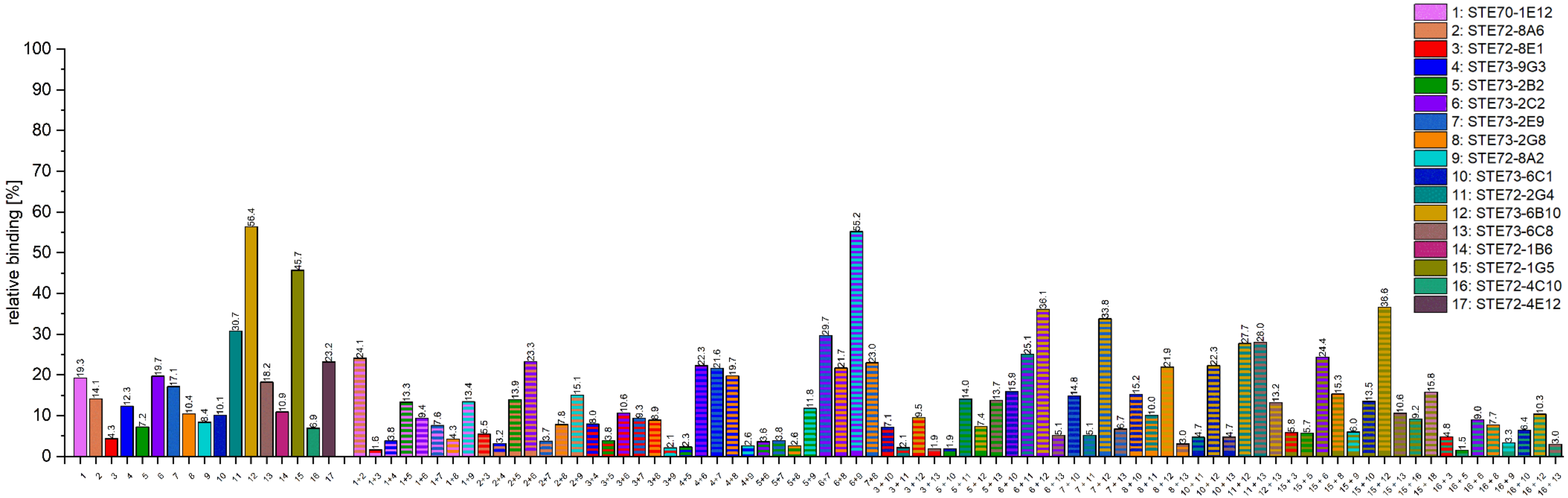

**Supplementary Data 6** Binding of the IgGs to RBD, S1 and S1-S2 in ELISA.

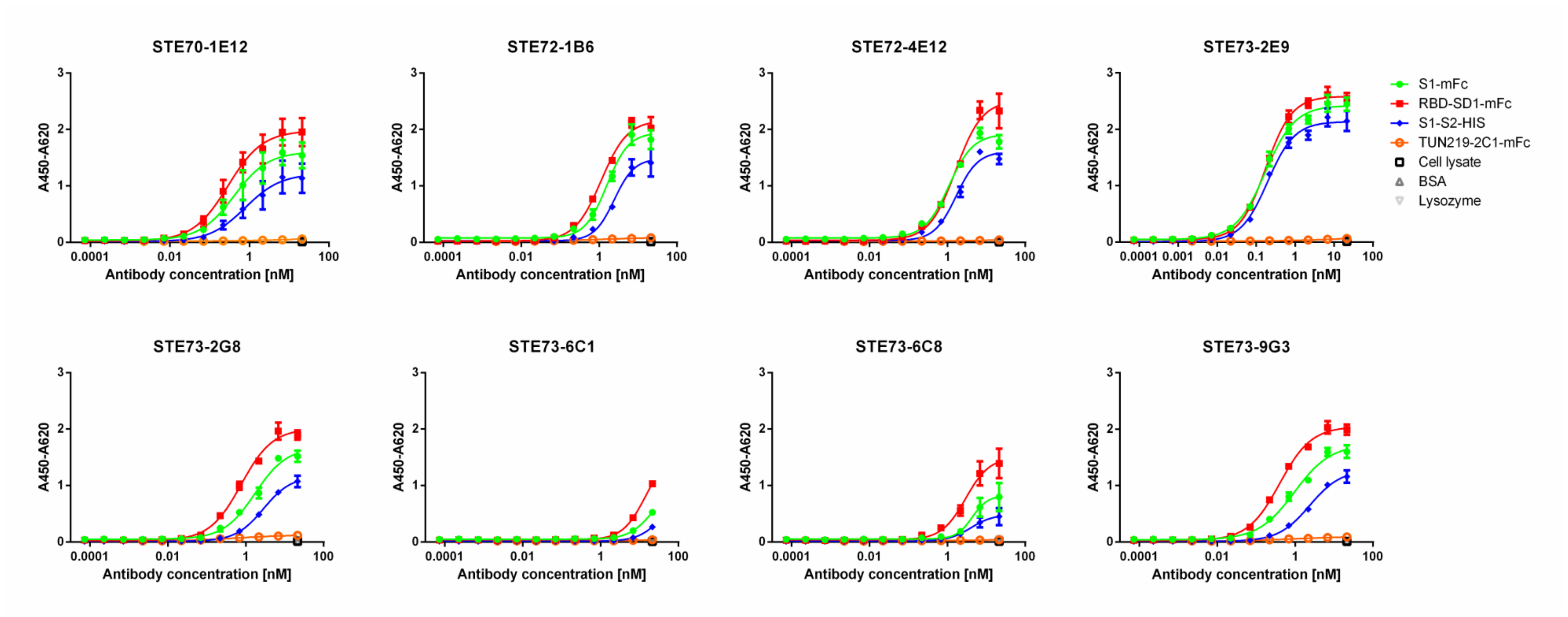

**Supplementary Data 7** Computational models of the antibodies (colored cartoon) on the spike trimer (surface, each monomer is in a different shade of grey). Only the variable region of the antibodies is shown.

**STE73-2E9**

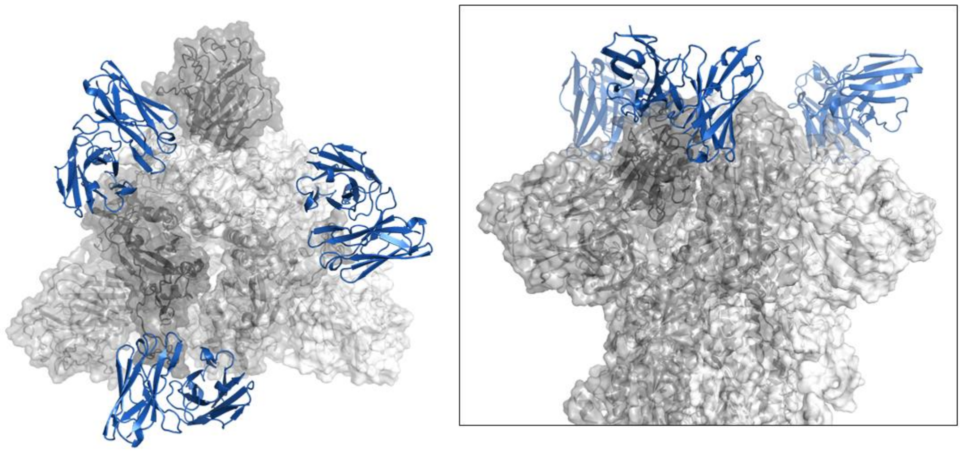

**STE73-2G8**

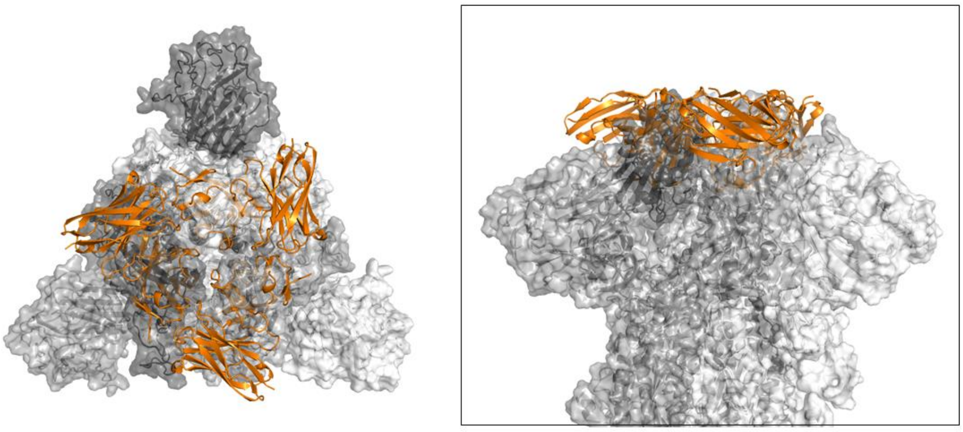

**STE73-9G3**

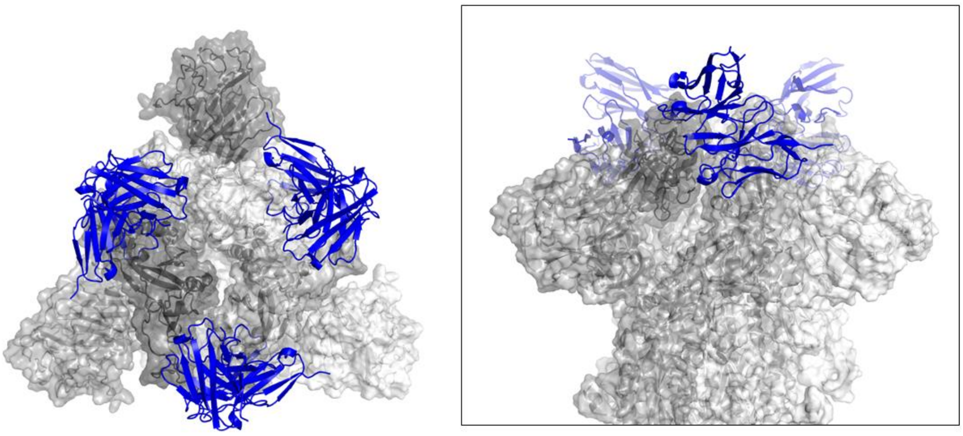

**Supplementary Data 8** SEC analysis of STE73-2E9 IgG under normal conditions (4°C, PBS, pH7.4), heat stress conditions (45°C for 24 h, PBS, pH7.4) and pH stress (4°C, 100 mM Na Acetate, pH3 for 24 h).

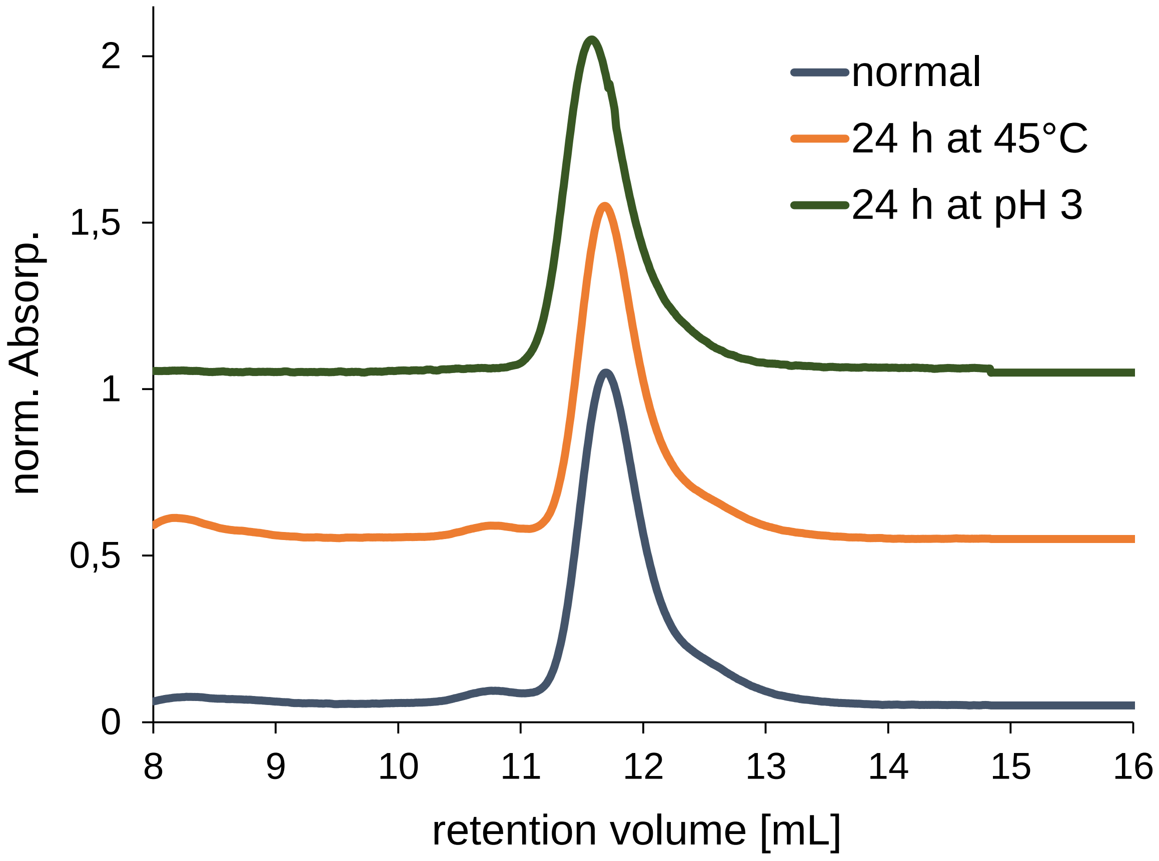
